# Utilising activity patterns of a complex biophysical network model to optimise intra-striatal deep brain stimulation

**DOI:** 10.1101/2024.04.12.589107

**Authors:** Konstantinos Spiliotis, Revathi Appali, Anna Karina Fontes Gomes, Jan Philipp Payonk, Simon Adrian, Ursula van Rienen, Jens Starke, Rüdiger Köhling

**Affiliations:** Institute of Mathematics, University of Rostock, Rostock, Germany; Institute of General Electrical Engineering, University of Rostock, Rostock, Germany; Faculty of Computer Science and Electrical Engineering, University of Rostock, Rostock, Germany; Oscar-Langendorff-Institute of Physiology Rostock University Medical Center Rostock, Germany

**Keywords:** Deep brain stimulation, Biophysical network model, Synchronisation

## Abstract

In this study, we develop a large-scale biophysical network model for the isolated striatal body to optimise potential intrastriatal deep brain stimulation applied in, e.g. obsessive-compulsive disorder by using spatiotemporal patterns produced by the network. The model uses modified Hodgkin-Huxley models on small-world connectivity, while the spatial information, i.e. the positions of neurons, is obtained from a detailed human atlas. The model produces neuronal activity patterns segregating healthy from pathological conditions. Three indices were used for the optimisation of stimulation protocols regarding stimulation frequency, amplitude and localisation: the mean activity of the entire network, the mean activity of the ventral striatal area (emerging as a defined community using modularity detection algorithms), and the frequency spectrum of the entire network activity. By minimising the deviation of the aforementioned indices from the normal state, we guide the optimisation of deep brain stimulation parameters regarding position, amplitude and frequency.

## Introduction

The striatum constitutes a subcortical region which loops information from the cortex via the other basal ganglia nuclei and the thalamus back to the cortex, thereby orchestrating such varied activities as motor control, decision-making, choosing actions, and, importantly, also reward behaviour [Calabresi et al. 2007, 2014], Crittenden and Graybiel [2011]. The striatum integrates cortical signals (prefrontal, motor, cerebral cortex) to create motor activities based on experience and forthcoming selections. Structurally, the caudate nucleus and the putamen form the striatal structure. On a cellular level, the striatum mainly consists of GABAergic medium spiny neurons (MSN) (95%), projecting mainly out of the striatum, with collaterals to other MSN, and interneurons (5%, most of these parvalbumin-positive) Assous and Tepper [2019], Straub et al. [2016], Gittis AH 2012].

The striatum has long been known to be involved in several neurological diseases resulting in movement disorders, ranging from Parkinson’s disease (PD) via Dystonias to Huntington’s disease (HD) as a widespread neurodegenerative disorder. Importantly, the striatum also impacts cognitive and reward processes (particularly the dorsal striatum) and hence, striatal function has been recognised to be pivotal in psychiatric conditions such as obsessive-compulsive disorder (OCD), depression, impulsivity, and attention-deficit hyperactivity disorder (ADHD) Crittenden and Graybiel [2011], Remijnse et al. [2006]. Deep brain stimulation of the striatum has therefore been introduced relatively recently as a novel approach for the treatment of OCD Graat et al. [2017], Widge et al. [2018, 2016]

It is long known that dopaminergic projections from the substantia nigra are important for striatal functionality, affecting both the direct and indirect pathways Albin et al. [1995], Gonon and Bloch [1997] (activating the former via D1 and dampening the latter via D2 receptors), and thus are crucially involved in the initiation of movements. A substantial part of the dopaminergic projections to the striatum (in particular the dorsal striatum), however, also comes from the ventral tegmental area (VTA)Kwon and Jang [2014], Derdeyn et al. [2022], which also projects to the nucleus accumbens, amygdala and prefrontal cortex. The dorsal striatum, in turn, projects to similar areas: the amygdala and the prefrontal cortex via the dorsal thalamic nuclei. In this way, the dorsal striatum has been recognised to be particularly involved in decision-making, goal-directed actions and reward mechanisms Balleine et al. [2007]. Consequently, low dopamine levels and disturbed striatal activity are linked with diseases involving movement disorders, but also depression and other neuropsychiatric diseasesCalabresi et al. [2014], Belujon and Grace [2017], Pizzagalli et al. [2019].

The main other input to the striatum is, obviously, cortical. This is provided as glutamatergic, excitatory cortico-striatal projections to medium spiny neurons Lassus et al. [2018]. Glutamatergic innervation to the striatum mainly projecting from cerebral and frontal cortex Paraskevopoulou et al. [2019] also sets the activity level of GABAergic neurons in the way of feed-forward inhibition and modulates striatal outputs controlling motor behaviour refer Fig.(1). Additionally, glutaminergic activation (through NMDA and AMPA receptors) regulates the formation of the synapses Paraskevopoulou et al. [2019], Chang et al. [2014].

Pathological conditions in the basal ganglia network are usually accompanied by changes in glutamatergic and dopaminergic signalling, suggesting that the dynamic interaction of these two inputs into the striatum becomes complex in disease. Postmortem studies in tissue from the caudate and putamen of patients with PD (compared to tissue from persons with causes of death unrelated to the brain) showed a reduction of about 27% in dendritic spines of MSNs Stephens et al. [2005]. Additionally, the MSN neurons in the caudate nucleus displayed a significant length reduction in PD. Dendritic spines receive crucial excitatory input from the cortex; thus, the spine density reduction in conjunction with the total dendritic length decrement was thought to have a negative impact on the excitatory tone of MSN neurons Stephens et al. [2005].

Similar conclusions can be drawn more generally from the classical model of basal ganglia activity Obeso and Lanciego [2011], Caravaggio et al. [2016], Albin et al. [1995]. Specifically, the classical model of the basal ganglia predicts that loss of striatal dopamine will decrease extracellular levels of glutamate in the striatum and cortex Albin et al. [1995], Obeso and Lanciego [2011]. Abnormalities in these dopaminergic and glutamatergic systems have been observed in numerous disorders, including Parkinson’s disease Loane and Politis [2011], and beyond movement disorders, also in depression, OCD and schizophrenia Musazzi et al. [2013], Howell et al. [2019]. Notably, previous fMRI studies Remijnse et al. [2006], Rao et al. [2010] in patients with OCD showed decreased responsiveness and activity in the ventral striatal and caudate. Conversely, healthy controls had increased perfusion in all striatal areas compared to the patients Remijnse et al. [2006].

Further studies suggest that the glutamatergic and dopaminergic input interact complexly beyond the basal ganglia Caravaggio et al. [2016], Scott et al. [2002], Chen et al. [2004], Howes and Kapur [2009]. Thus, it has been shown that NMDA receptor activation regulates D1-dopamine receptor signalling in cortical neurons and vice versa Scott et al. [2002], Chen et al. [2004], Howes and Kapur [2009]. Specifically, activation of D1-receptors in the prefrontal cortical pyramidal neurons by the agonist SKF81297 increases the stationary NMDA evoked current Chen et al. [2004]. Inversely, activation of NMDA receptors by glutamate results in the recruitment of D1-receptors in cortical neurons, while they do not affect the distribution of D2-receptors Scott et al. [2002]. Taking these findings together, we hypothesise that any lesion of the glutaminergic-dopaminergic circuit leads to malfunctioning and, consequently, reduced striatal activity.

Deep brain stimulation (DBS) of the striatum has evolved as a promising therapy for patients with severe and resistant forms of OCD and mental impairments Blomstedt et al. [2013], Wu et al. [2021], Widge et al. [2019]. As in the case of DBS for PD, considerable unknowns remain, including the anatomical targets of stimulation, optimal stimulation parameters, long-term effects of stimulation, and the patient’s clinical and biological response to DBS. Progress in predicting therapeutic DBS effects (by optimizing DBS parameters: position, intensity, frequency etc.) was achieved using different computational models Butson et al. [2007], Butenko et al. [2020], Rubin and Terman [2004], Popovych and Tass [2019], Spiliotis et al. [2022], Fleming et al. [2020]. However, due to the strongly heterogeneous nature of the connection topology and intrinsic complexity (stochastic and nonlinear large scale neuron, multiple scales) Spiliotis and Siettos [2011], Siettos and Starke [2016, Deco et al. 2008, 2013], Bassett and Bullmore [2017], Iliopoulos and Papasotiriou [2022], DBS outcome is far from trivial to predict.

Towards this direction, we extend a previous model of the striatal activity Chartove et al. [2020], completing the DBS action in the mathematical model. Specifically, the contribution of this work is two-fold. (i) It presents a large-scale biophysical model of the striatum to predict neural activity and spatial-temporal patterns, and, using the model, (ii) it optimises DBS parameters (namely, placement of electrodes, frequency, and amplitude).

We obtain (i) by modelling the neurons with modified Hodgkin-Huxley equations Chartove et al. [2020], Hodgkin and Huxley [1952] and using a complex graph structure to model the neural network. We construct the graph structure from the real coordinates of human atlas Iacono et al. [2015], placing two types of striatal neurons: interneurons as fast-spiking (FS) and MSN neurons. Going beyond other studies published before Chartove et al. [2020], we use complex connectivity (i.e., small world structures Watts and Strogatz [1998]) simulating more realistic neuron connectivity structures Bassett and Bullmore [2017, 2006]. Integrating this high-dimensional nonlinear system, we produce spatiotemporal patterns of striatal activity. Our model predicts, similar to the classical model of abnormal activity of basal ganglia Obeso and Lanciego [2011], Caravaggio et al. [2016], reduced striatal activity when a cortico-striatal connectivity breakdown is simulated, which renders it a candidate for deriving optimised DBS parameters. (ii) To obtain optimised DBS parameters, we use three different macroscopic measures or biomarkers: (a) The mean network activity of the striatum (b) a combination of the mean activity with the rhythmicity of the network(i.e., the spectrum of mean membrane potential of neurons, using Fourier analysis), and (c) the mean activity restricted to a specific area or community of striatum, that is, the dorsal striatum, offering a more targeted treatment. Based on the deviation of the indices from the healthy state, we propose a DBS protocol that provides the therapeutic pattern for abnormal striatum network activity. Unlike other models, we include a sensitivity analysis gauging the impact of specific parameters on total network activity and corroborating our assumption that individual parameters play a paramount role.

## Materials and methods

### Data sources

The structural Magnetic Resonance Imaging (MRI) data are taken from the Human Connectome Project (HCP) VAN 2012]. These MRI data were laid over another atlas, the MNIPD25 atlas, which is most commonly used for surgery planning Xiao et al. [2017, 2012], CHA 2006] with the help of Lead-DBS, a toolbox for atlas connectivity and segregation analyses Horn et al. [2019]. The structural composition striatum, in turn, is taken from the MIDA atlas Iacono et al. [2015] and translated into the MNI (Montreal Neurological Institute) space by using the segmented MRI data as a reference.

### Modelling intrastriatal connectivity using complex networks

In contrast to other models of the striatum, we leverage a graph network modelling the connections of neurons under the hypothesis the graph properties are linked to biological activities. Thus, four main graph indices characterise the structural connectivity of the network, i.e. degree distribution (number of connections of each neuron), connection efficacy (minimal distance between nodes), clustering coefficients (numbers of locally interconnected triplets) and the degree of betweenness-centrality, i.e. the number of neurons serving as high-density hubs. The knowledge of the topological structure of the network plays an important role in emergent neural activity and functionality Deco et al. [2012, 2008]. Furthermore, knowledge of structural details allows us to investigate the mechanisms involved in neural functionality or dysfunctionality. Network structural properties, in turn, can be identified using network statistical measures as listed above (i.e. degree distribution average path length, clustering coefficients, centrality Stam and Reijneveld [2007], Watts and Strogatz [1998], Spiliotis et al. [2022]). In our case, the positions of nodes were defined by the data source analysis. How these nodes are connected to each other will be described in the following section.

### Construction of complex network using the structural connectivity

Initially, data source analysis defined a network consisting of 1,995 nodes. In this network, we assume that the majority of the nodes (95%) represent MSN neurons, and the remaining 5% are interneurons, consistent with Yager et al. [2015]. The connectivity of the striatum is constructed following the idea of the small-world algorithm Watts and Strogatz [1998], Bassett and Bullmore [2006], Bullmore and Sporns [2009], Stam and Reijneveld [2007], Spiliotis and Siettos [2011]: initially, each MSN neuron is connected with *k* = 20 neurons in the vicinity of 5 mm (local connections). Next, with a small probability *p*, and for each local connection, a new remote neighbour is added. For interneurons, we follow the same structure, using, however, a five times denser interneuron to MSN connectivity (i.e. *k* = 100). In this way, the resulting network is highly clustered (like a lattice structure), additionally with a small distance between nodes (like random networks).

Unlike traditionally used lattice connectivity models or all-to-all models, small-world structures better represent the physiological networks as a result of two main characteristics Netoff et al. [2004], Berman et al. [2016], She et al. [2016], Bassett and Bullmore [2006, 2017], Fang et al. [2017], de Santos-Sierra et al. [2014]: they are highly clustered, and typically show short path lengths Bullmore and Sporns [2009], Watts and Strogatz [1998], Newman [2006], Spiliotis et al. [2022], enhancing in this way signal or rhythm propagation within the network, and the synchronizability of the network. The resulting striatal network is represented as graph *G* = (*V, E*), where *V* is the set of nodes and *E* represents the set of edges, i.e. connections. The nodes of the structural network are defined as points in three-dimensional space, as a single neuron with spatial coordinates. The connectivity can be represented with the adjacency (or connectivity) matrix *A*: if two neurons at positions **x** = (*x*_1_, *y*_1_, *z*_1_) and **y** = (*x*_2_, *y*_2_, *z*_2_) are linked then *A*(**x, y**) = 1, otherwise *A*(**x, y**) = 0. In the next section, we provide tools that allow us to extract the connectivity properties of the striatal network.

### Network measures characterising striatal connectivity

Network measures Bullmore and Sporns [2009] reveal important properties of the structure, e.g. the expected number of connections for each node of the network, the centrality of nodes (importance of nodes), the average shortest path (expected number of steps between any two nodes), and, importantly, the detection of communities (or modules) and how these organise the network by creating dynamical patterns Deco et al. [2012, 2008]. A network measure can express a specific property of a node (e.g. the clustering property of one node) and appears as an individual property of the *i*th node, while the resulting distribution over the whole network defines the global-macroscopic description (e.g. the statistical distribution of clustering).

### Degree distribution

The degree of a node *i* refers to the number of links connected to it Bullmore and Sporns [2009]. In directed networks, a node has both an in-degree and out-degree, numbering in-coming and out-going edges, respectively. A high degree of connectivity (increased numbers of links) of the *i*th node defines the importance of a node in the network. The degree distribution *P* (*k*) defines the probability of a randomly selected node having a specific degree *k*. Fig. 2A shows the degree distribution for the striatal network. The probability density can be separated into two parts: one corresponds to MSN neurons with a mean degree of around 30 and a second with a mean degree of 100, corresponding to interneurons. Both peaks show a symmetry around each mean value (e.g. 30 and 100).

**Fig 1:**
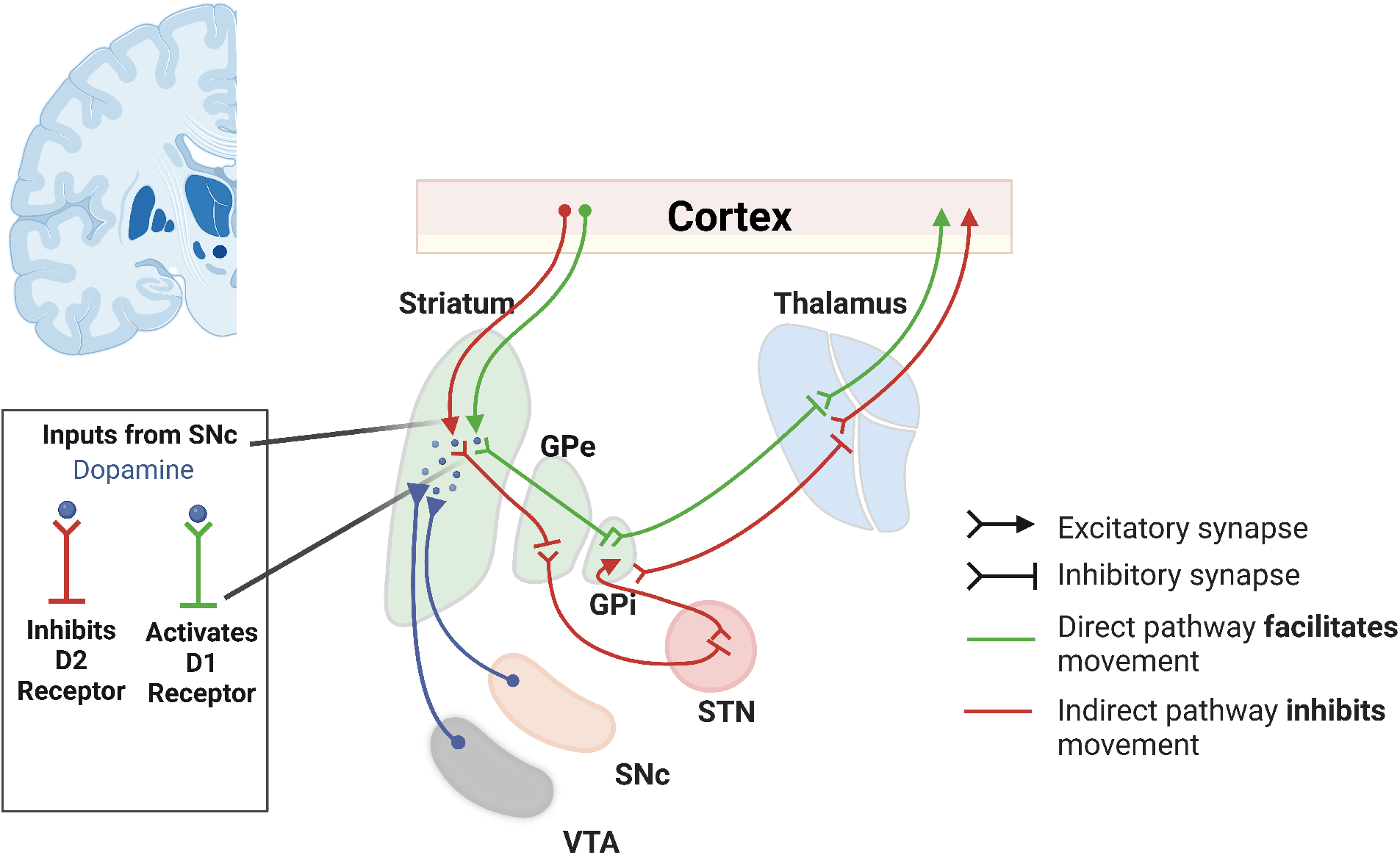
Pathways and circuits of basal ganglia. Direct and indirect basal ganglia pathways are depicted with inputs from the substantia nigra. Created with biorender.com

**Fig 2:**
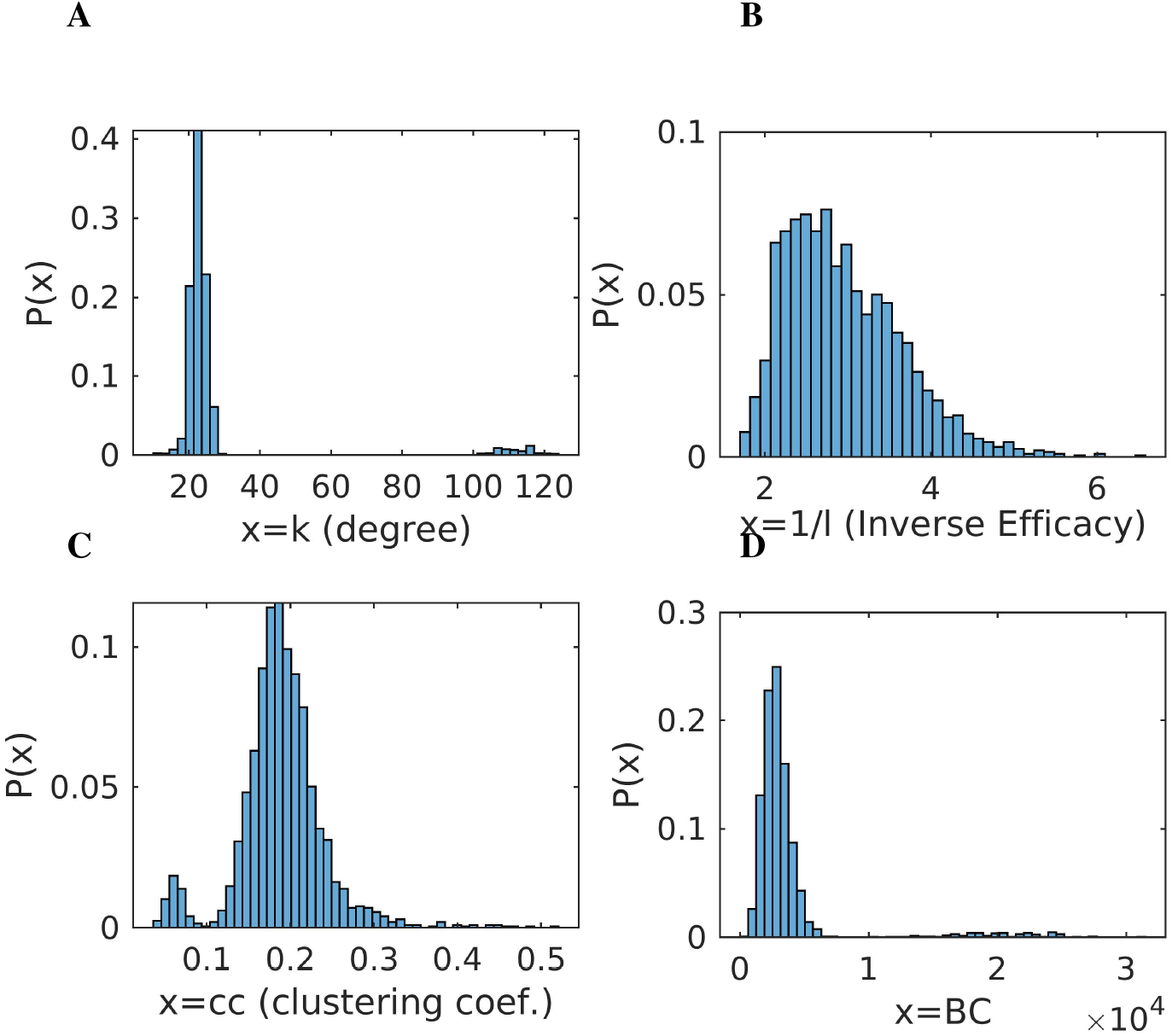
Connectivity properties of the striatal network. **A** Distribution of the number of connections (*Degree distribution*) of any one neuron. The distribution reflects the presence of two cell types, each showing a relatively symmetrical spread of data: medium spiny neurons (the majority) with relatively sparse connections (mean 22 connections) and fast-spiking GABAergic interneurons (minority; mean 110 connections). **B**. Distribution of connection *efficacy* (the number of nodes as maximal distance between any two neurons, here given as reciprocal value to avoid divisions by 0). **C** The distribution of *clustering coefficients*, i.e. the number of locally interconnected neuronal triplets. **D** The distribution of *betweenness centrality* (BC), measured as the number of possible paths converging onto any given neuron. Again, the distribution is double-peaked, with approx. four thousand paths converging onto the majority of neurons, while some neurons have 10,000 to 20,000 paths converging onto them.

### Path lengths, efficacy and clustering coefficient

The minimum number of steps between two nodes in the network (in the case of binary networks such as the one described here) defines the shortest path length *d*_*i→j*_ between node *i* and *j*. Averaging over the set of all shortest paths, we obtain the mean path length of the network Bullmore and Sporns [2009]:

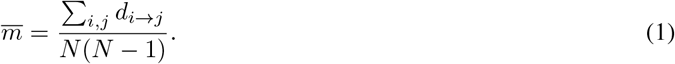

The mean path length shows the ability of the network to spread information (signal activity) between any two nodes. A low mean shortest path length 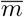 highlights that any two randomly chosen nodes can interchange information very fast (passing through very few intermediate nodes; in our case ≈ 4 nodes).

If no pathway exists between node *i* and *j* then *d*_*i→j*_ = ∞, and this pair of nodes is excluded from eq. (1). A similar measure reflecting pathway length, which, however avoids divisions by 0, is the global efficacy 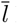 Bullmore and Sporns [2009]:

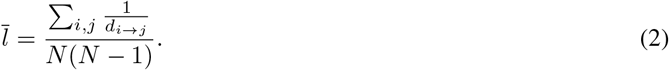

now *d*_*i→j*_ = ∞ =⇒ 1*/d*_*i→j*_ = 0.

The inverse global efficacy 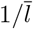 is thus reflecting the mean shortest path. For the striatal network, the inverse global efficacy is computed 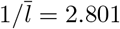. Fig. 2B shows the distribution of distances between any two nodes.

Another important measure which characterises the connectivity is the clustering coefficient. It measures the proportion of triangle loops that exist in a node, expressing a feedback mechanism that enhances the rhythm generation. Specifically, the clustering coefficient of a node *i* is defined as ratio:

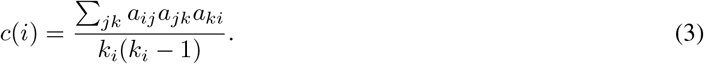

The higher the number of triangles (that exist) with respect to the *i*th node, the higher the clustering coefficient. Fig. 2C depicts the distribution of clustering coefficients. The mean clustering coefficient is computed as 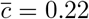. The distribution shows the existence of a few nodes with high values of *c*.

### Betweenness centrality

Another important measure which quantifies the significance of a node is ‘betweenness centrality’. The term ‘centrality’ is related to the degree of influence which this particular node exerts within the network. ‘Betweenness centrality’ thus measures the amount of influence which a node has with respect to the total information flow in the network. Important nodes that connect different subgraphs in the network (i.e., act as a bridge) show high betweenness centrality. The betweenness centrality *b*_*c*_(*i*) of the *i*th node is mathematically defined as the fraction of all shortest paths in the network that pass through the node, that is,

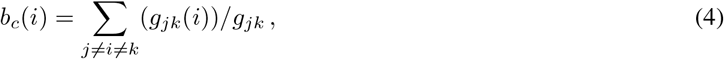

where *g*_*jk*_(*i*) is the number of shortest paths from *j* to *k* passing over *i*, and *g*_*jk*_ is the number of shortest paths between nodes *j* and *k*. Bridging nodes that connect different subsets of the network often have a high betweenness centrality. Higher values of *b*_*c*_(*i*) indicate that the node acts as a central hub. The importance of these hubs is also highlighted pathophysiologically as such hubs are ideally suited as targets of therapeutic intervention, i.e. for DBS Spiliotis et al. [2022].

Fig. 2D depicts the distribution of *b*_*c*_ of the network. As becomes evident, the large majority of nodes show very low centrality. However, there are few nodes with high *b*_*c*_. In Fig. 3, black-filled circles depict the spatial localisation of these central nodes in the network.

**Fig 3:**
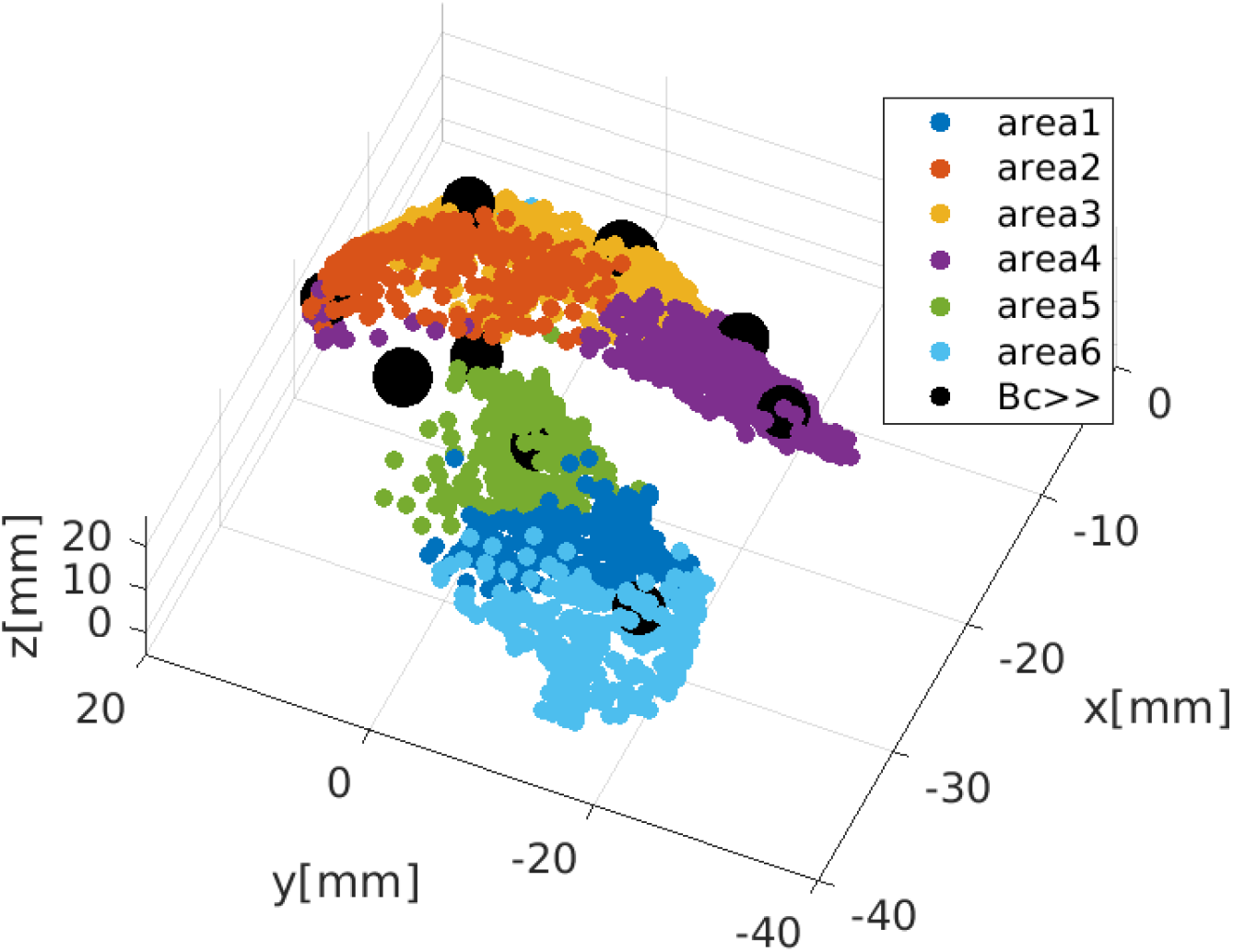
Modularity: Community detection in the striatal network. Communities were extracted using an iterative process of branching groups of neurons fulfilling two criteria: a) dense connectivity among members, and b) sparse connectivity to the other communities. This iterative branching stops when an optimum is reached in any branch. In this way, the striatal network is partitioned into separate subgraphs, using a commonly used modularity-index algorithm Leicht and Newman [2008]. Using a boundary condition that at least 180 neurons should be within one community (9 % of the population), in the present model, the algorithm identifies six communities, which remain stable with repetitive (20 times) realisations. As the figure shows, three communities are located in the caudate nucleus (top, red, yellow and violet hues) and three in the putamen (bottom, blue and green hues). The ten nodes with the highest betweenness-centrality (hubs, black circles) are equally distributed between the caudate nucleus (n = 5) and the putamen (n = 5).

### Detection of communities and modularity

Networks characteristically contain subsets (subgraphs) that have dense internal connectivity (connectivity among nodes in the subset) and sparse connections to other subgraphs Newman [2006]. The partition of the network into densely connected subgraphs (or communities) plays a significant role in information processing within the network, and it is also related to different biological functions of the area (e.g. striatum) Voorn et al. [2004]. Assigning and allocating these densely connected communities to brain structures allows to construct a modular view of the dynamics of the network Mylonas et al. [2016], Betzel et al. [2017].

The modality index identifies such densely connected communities. The modality index Newman [2006] assigns a community number *s*_*i*_ to each node. For example, if there are two communities, then *s*_*i*_ = ± 1. Here, we seek the best network partition in order to maximise the modularity function *Q*:

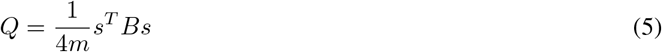

where *m* = 1*/*2 Σ*k*_*ij*_ is the total number of edges in the network, and *B*_*ij*_ = *A*_*ij*_ − *k*_*i*_*k*_*j*_*/*2*m* is the resultant modularity matrix, also known as graph Laplacian matrix. In such matrices, the optimisations can be achieved using graph partitioning or spectral partitioning (eigenvalues-eigenvectors decomposition) of *B* Newman [2006], Leicht and Newman [2008].

Figure 3 shows the communities for the striatal network as determined by the optimisation of the *Q* function. The detected communities emerge as positioned within the boundaries of the brain nuclei; i.e., they follow striatal anatomy, which is likely related to functional somatotopy in the sense that ‘the limbic loop connecting the ventral striatum with the ventromedial prefrontal cortex (vmPFC) has been implicated in motivational and emotional processing, whereas the associative and sensorimotor networks regulate different forms of behaviours in instrumental behaviour processing. The associative network connecting the dorsomedial striatum with the dorsolateral prefrontal cortex (dlPFC) mainly contributes to goal-directed behaviours, while the sensorimotor network projecting from the dorsolateral striatum to the sensorimotor cortex is mainly responsible for the habitual control behaviours in instrumental learning’ Voorn et al. [2004]. In our case, six communities emerged from the simulation as populations with 294, 473, 189, 399, 290, and 330 members, respectively.

### Modelling and simulation of MSN networks Modelling MSN neurons

The dynamics of each MSN neuron are modelled by current balance equations for the membrane potential: Chartove et al. [2020]

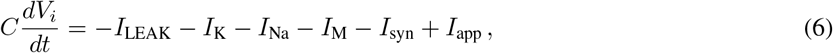

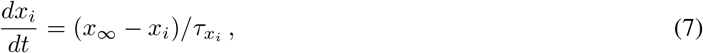

where *C* is the membrane capacity, and *V*_*i*_ is the membrane potential of the *i*th neuron. The current balance eq. (6) contains four membrane currents Chartove et al. [2020], the fast sodium and potassium currents *I*_Na_ and *I*_K_, the leak current *I*_LEAK_, and an M-current *I*_M_. All ionic currents follow the Hodgkin-Huxley formalism Hodgkin and Huxley [1952]: 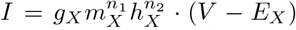, where the exponents *n*_1_, *n*_2_ represent the number of activation-inactivation channels, respectively, *g*_*X*_ is the maximum conductance of the *X* ion, and *E*_*X*_ stands for the reversal potential for each *X* ion (*X* ∈{Na, K}). Specifically, the sodium current has three activation gates and one inactivation gate, that is,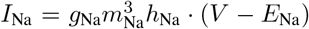. The potassium current has the form 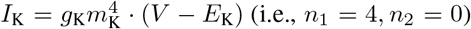 (i.e., *n*_1_ = 4, *n*_2_ = 0). Finally, the M-current and leak currents follow *I*_M_ = *g*_M_*m*_M_ · (*V* − *E*_K_) and *I*_LEAK_ = *g*_LEAK_ · (*V* − *E*_LEAK_), respectively.

The variable *x*_*i*_ denotes the gating variables *m*_X_ and *h*_X_. Following the Hodgkin-Huxley formalism Hodgkin and Huxley [1952], Chartove et al. [2020], the function *x*_*∞*_ is given by 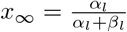 and the time by 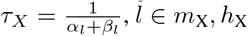, *l* ∈ *m*_X_, *h*_X_. For the sodium current and for the gating variable *m*_X_, we obtain

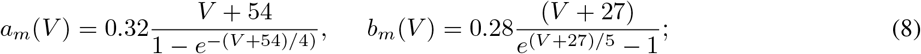

similarly, for the gating variable *h*_X_:

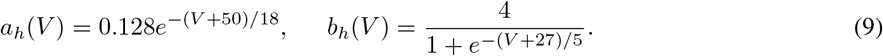

For the potassium current, with only one activation gating: *m*_k_

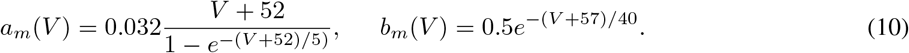

For the M-current, we obtain

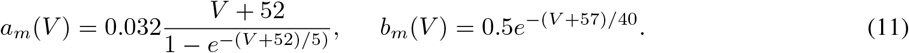

The current *I*_app_ is written as *I*_app_ = *I*_0_ + *I*_DBS_ in eq. (7), where *I*_0_ represents a network activation current, describing the dependence of the neuronal activation due to dopamine receptor activation, or due to intensity of cortical-striatal connectivity. The current *I*_DBS_ specifically models deep brain stimulation, and it is applied on any element within the reach of the stimulation electrodes (the stimulation is modelled as declining exponentially, see eq.(12)). The mathematical description is given by the form:

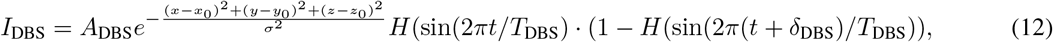

while in the absence of DBS treatment *I*_DBS_ = 0.

The synaptic currents *I*_syn_ for MSN neurons can be written as a sum: *I*_syn_ = *I*_MSN *→* MSN_ + *I*_FS*→* MSN_, where *I*_MSN*→* MSN_ is the inhibitory synaptic current between MSN neurons, carried by fast-spiking neurons, whose exact description is given in the next section.

**Table 1:**
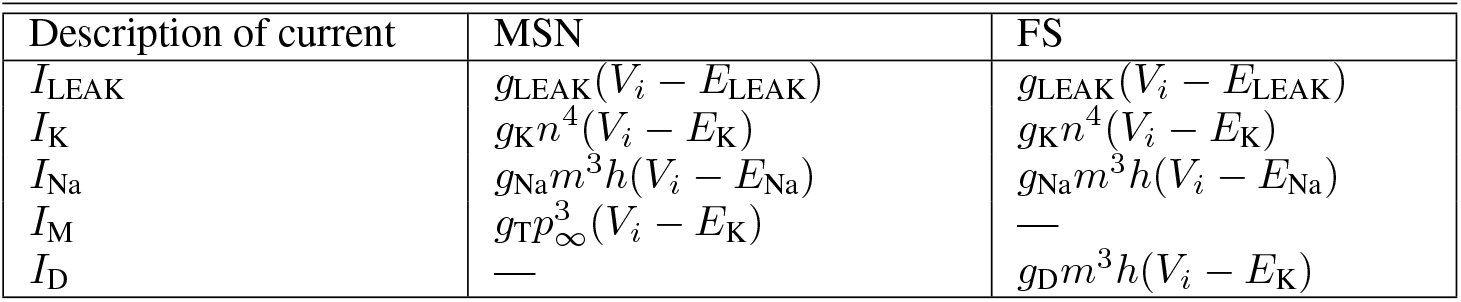
The currents for MSN and fast-spiking neurons.

### Modelling Fast Spiking (FS) neurons

The dynamics of each FS neuron are modelled by current balance equations for the membrane potential: Chartove et al. [2020]:

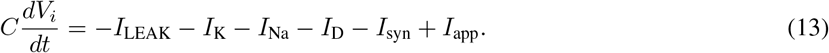

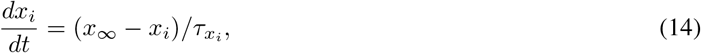

where *C* is the membrane capacity, and *V*_*i*_ is the membrane potential of the *i*th FS neuron. The current balance eq. (14) contains four membrane currents Chartove et al. [2020]: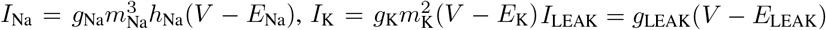, *I*_LEAK_ = *g*_LEAK_(*V* − *E*_LEAK_), while the fast-activating, slowly inactivating dendritic potassium D-current has the form Chartove et al. [2020], Golomb et al. [2007]: 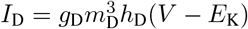, with three activation gates and one inactivation gate (i.e. *n*_1_ = 3, *n*_2_ = 1), thus imposing a delay in firing upon depolarization. For the sodium current *I*_Na_, and for the gating variable *m*, we obtain:

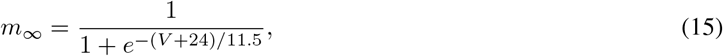

and for the sodium inactivation

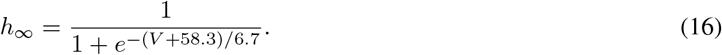

For the potassium *I*_K_current, and for the activation *m* variable, we use the equation:

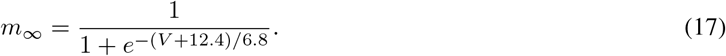

Finally, for the D-current with activation and inactivation variables, we use

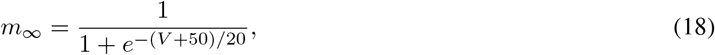

and for the inactivation

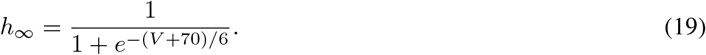

### Description of the network inhibitory synaptic activity

The coupling between the neurons in eqns. (6) and (13) is described by the synaptic current *I*_syn_. Initially, we model the activation of a synapse using the activation variable *s*_*i*_ (for the *i*th neuron), which is given by Laing and Chow [2002], Ermentrout and Terman [2012], Compte et al. [2000]:

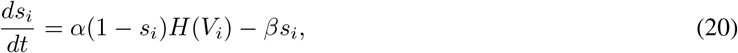

where the function *H*(*V*) is a smooth approximation of the step (Heaviside) function *H*_step_ (i.e. *H*_step_(*x*) = 1, *x >* 0 and *H*_step_(*x*) = 0, *x <* 0.) The variable *s*_*i*_ describes the activation of synapses from the pre-synaptic neuron *i* to the post-synaptic neuron *j*. The parameters *α, β* in eq. (20) are related to the activation and inactivation time scales, respectively, of the inhibitory (GABA-ergic) synaptic connections. In cases of MSN - MSN and MSN - FSI interactions, the form of the function *H* is Chartove et al. [2020]:

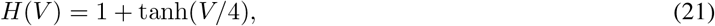

with activation rates in equation (20) *α* = 2, *β* = 1*/*13 ≈ 0.08. Similarly, for FS - MSN and FS - FS interactions, the form of the function *H* is given as:

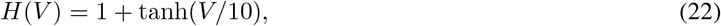

with activation rates in equation (20) *α* = 4, *β* = 1*/*13 ≈ 0.08.

For each *i* th neuron in the network, the total synaptic inhibition which it receives from the pre-synaptic neurons is:

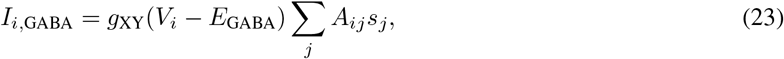

with *E*_GABA_ = − 80*mV*. The matrix element *A*_*ij*_ has the value 1 or 0, depending on whether neurons *i* and *j* are connected or not. In this way, it resembles the modified Watts and Strogatz (WS) small-world topology Watts and Strogatz [1998], Bullmore and Sporns [2009], Stam and Reijneveld [2007], Gafarov [2016], De Santos-Sierra et al. [2014], Netoff et al. [2004], Bertalan et al. [2017], Fang et al. [2017]. The summation is taken over all presynaptic neurons. The parameter *g*_XY_ represents the conductance between *X* and *Y* interactions *X, Y* ∈ {MSN, FS}.

### Modelling the connectivity within the striatum

The synaptic current for the MSN neurons is given by:

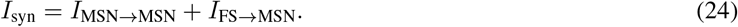

The current *I*_MSN_ *→* _MSN_ indicates the inhibition between MSN-MSN neurons, while the second term *I*_FS_ *→* _MSN_ represents the interneuronal inhibition. Taken together, the mathematical form of the synaptic current for MSN neurons is:

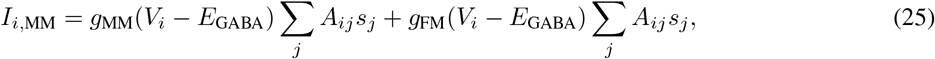

where, again, the element *A*_*ij*_ has the value 1 or 0, depending on whether neurons *i* and *j* are connected or not, while the sum is taken over all presynaptic neurons.

Similarly, for the FS neurons, the synaptic current is analysed as a sum:

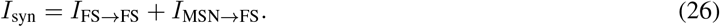

The current *I*_FS_ *→* _FS_ represents the rare case of FS-FS inhibition, while the second term *I*_MSN_ *→* _FS_ imitates the feedback inhibitory loop of MSN to interneurons. Then, the mathematical form of the synaptic current for each FS neuron is:

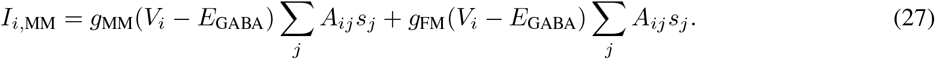

### Restoring normal striatal activity by optimising DBS position

As already emphasised, striatal neuronal activity is not only involved in major tasks such as movement control but also in decision-making, reward behaviour and other cognitive/emotional tasks, with behavioural control being driven by the ventral parts of the striatum Calabresi et al. [2007, 2014]. Under pathological conditions (e.g. obsessive-compulsive disorder), abnormal striatal activity has been reported; decreased dopamine levels in conjunction with aberrant cortico-striatal interactions are thought to lead to reduced striatal network activity Chartove et al. [2020], Rao et al. [2010].

In order to quantify striatal activity, we define the mean network activity as a macroscopic variable Gerstner et al. [2014]:

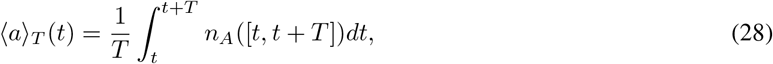

where *N* is the number of neurons in the population, *n*_*A*_([*t, t* + *T* ]) is the number of spikes (summed over all neurons in the population) that occur between *t* and *t* + *T*, and where T is a small macroscopic time interval (10 ms). The values obtained were then divided by the number of bins (100); thus, ⟨*a*⟩ _*T*_ actually stands for the number of spikes per 0.1 ms within the entire population of neurons (2000 neurons), and we define ⟨*a*⟩ _*T*_ as the mean network activation rate. Low values of the macroscopic network rate indicate low striatal output, characteristic of a disturbed and abnormally low dopamine, or low intensity of corticostriatal activity.

Another macroscopic variable that we explore is the mean membrane voltage 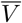 of neurons in the network; specifically, we define:

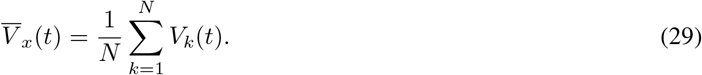

The mean voltage activity 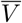 (indirectly related to the local field potential (LFP) in the case of supra-threshold values resulting in spiking) is utilised for the characterisation of synchronised rhythm (through Fourier spectrum) under different states (healthy, abnormal or abnormal plus DBS).

### Optimising DBS parameters using macroscopic quantities of the striatal network

Differences in striatal targeting areas, as well as different intensity and frequency values of the DBS signal result in differences in distant network activation. In the model, we vary position, stimulation intensity and frequency, resulting in the parameter vector *r* = (*x*_0_, *y*_0_, *z*_0_, *A*_DBS_, *f*) ∈ ℝ^5^ to estimate optimal DBS outcome; see eq. (12). The effectiveness of DBS is evaluated using three objective functions. The objective functions defined below indicate the impact of DBS, i.e. the ability of DBS to restore neuronal activity to a healthy state.

### Optimizing with respect to firing rate

The first objective function is based on the mean network activity *< a >*_*T*_ eq. (28). We define the objective function as the difference between healthy and DBS mean activities, i.e.

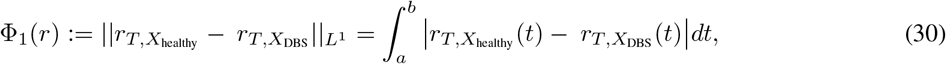

where *a, b* are the times of activation and inactivation of DBS (usually *a* = 0 and *b* = 1 sec), *r* = (*x*_0_, *y*_0_, *z*_0_, *A*_DBS_, *f*) ∈ ℝ^5^ is the DBS parameter vector, i.e. the position *r* = (*x*_0_, *y*_0_, *z*0) of the DBS electrode, the amplitude *A*_DBS_ and the frequency *f* of the pulse. The values of the model parameters, *r* ∈ ℝ^5^ were estimated numerically by minimising the residual, i.e.:

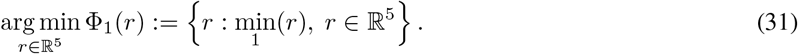

Minima of the difference functions Φ_1_, Φ_2_ and Φ_3_, i.e. values of the objective functions close to 0 imply 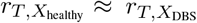, which it is interpreted as effective DBS action restoration of normal striatal activity).

### Optimizing with respect to network rhythmicity

The second objective function Φ_2_ is based on the frequency spectrum produced by the model. For this, we define the Fourier transform of the mean activity 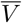 i.e.

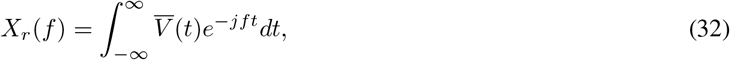

where *j* is the imaginary unit. The power spectrum is defined as |*P* (*f*) | = |*X*(*f*) | ^2^. To estimate the effectiveness of DBS, we calculate differences curves based on the following subtraction pairs: |*P*_*X*_ (*f*)| − |*P*_*Y*_ (*f*) |, where |*P*_*X*_ |, |*P*_*Y*_| are the power spectra of the states, *X, Y*, respectively. *X, Y* in this case represent the conditions under DBS *X*, and healthy state *Y*.

The objecting function Φ_2_ is then defined as the area under the curve (AUC, Valor et al. [2019]) for each of these difference pairs, i.e.

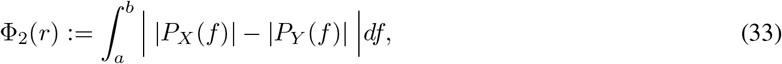

where *a, b* are the frequency range (i.e. a,b]=[0 300]Hz) and *r* = (*x*_0_, *y*_0_, *z*_0_, *A*_DBS_, *f*) ∈ ℝ^5^ is the DBS parameter vector. Similar to the previous case, i.e. Φ_1_, the values of the model parameter, *r* ∈ ℝ^5^ were estimated numerically by minimising the residual, that is,

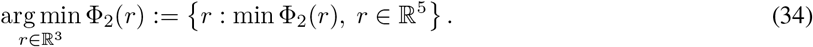

We can also combine the aforementioned objective functions to obtain a third optimisation scheme which takes into account both two macroscopic characteristics (i.e. rate and rhythmicity) of the network activity. We will present this scheme in the next subsection.

### Optimizing using a combination of network rhythmicity and firing rate

We define a combination of objectives functions eqs. (33) and (30):

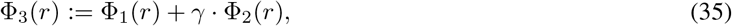

where *γ* is a scaling factor that balances (reduces or increases) the importance of the phase spectrum in the optimisation process. In this instance, *γ* is adapted by iterative optimisation steps with decreasing sizes of *γ*. At the chosen value, the objective function Φ_3_ combines both Φ_1_, Φ_2_ optimally, i.e. considers both the rate and the rhythmicity effect.

The minimisation problem was solved in all cases using a nonlinear least-squares solver (the MATLAB function **lsqnonln**). The step size tolerance was set to tol(*X*) = 0.001 and the function tolerance was set to tolF = 0.1. The maximum number of iterations was set to 100.

The last part of our theoretical analysis (Sensitivity Analysis) is a further investigation of the model parameters. Specifically, the aim is to examine the significance or the sensitivity of the emergent network behaviour with respect to important parameters such as the intensity of cortico-striatal connectivity and the connectivity conductance between MSN neurons. Using Sobol’ index as an order parameter i.e. Sobol ∈ [0, 1] (Sobol’ index ≈ 1 shows a greater influence of the parameter on the response of the model), we partitioning the variance of the output into fractions according to the parameter input’s contribution.

### Sensitivity analysis

For the global sensitivity analysis, i.e., to quantify the effects of the input random variables in the variance of the response of the model, we use Sobol’ indices Sobol [2001], based on functional decomposition applied to the variance. The implementation and use of the method are straightforward. In the system of interest, we consider **X** as a random input vector following a certain probability density function *f*_**X**_, and **Y** as the response of this system.

The total variance of a model’s response is denoted by var(**Y**), and the conditional variances, which consider the contribution of one or more parameters, are denoted by var[𝔼 (**Y** | **X**_*k*_)], where **X**_*k*_ is the input vector with *k* the parameters, with *i, k* ∈ {1, · · ·, *n* }. From Sobol’ decomposition and its orthogonality, we obtain the total variance of **Y** as the sum of the conditional variances, i.e.,

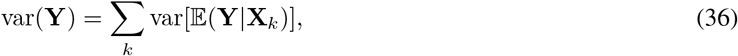

where 𝔼 is the expectation. In this sense, Sobol’ indices are defined by

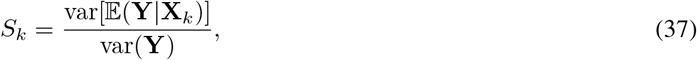

such that their total sum equals one. Therefore, a Sobol’ index can have a value within the interval [0,1]. The closer the sensitivity index of a parameter approaches the value 1, the greater its influence on the response of the model.

The aim is to examine the significance of a particular variable by measuring the proportion of variance in the Quantity of Interest (QoI) for which it is responsible. For this, we compute the first-order Sobol’ index, which quantifies the share of variance in the output due to the examined parameter. In addition, the higher-order index quantification considers the interaction of all studied variables.

To compute Sobol’ indices, we choose the Polynomial Chaos Expansion (PCE), which has proven to be very efficient in dealing with uncertainties. The PCE method is an efficient alternative to the Monte-Carlo methods, being much faster in obtaining similar results, as long as the number of uncertain variables is less than 20, see, e.g., Crestaux et al. [2009]. PCE can significantly decrease the number of simulations while providing an accurate approximation of the model’s response.

The PCE method uses an approximation of a system’s random model response **Y**, assuming that **Y** has finite variance.

The representation of **Y** can be given by

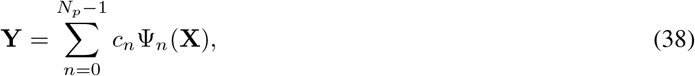

with *N*_*p*_ as the number of expansion factors, *c*_*n*_ as the coefficients, and Ψ_*n*_ as a multidimensional generalized PC basis defined in a Hilbert space *L*^2^(**X**, *f*_**X**_). The coefficients are computed following

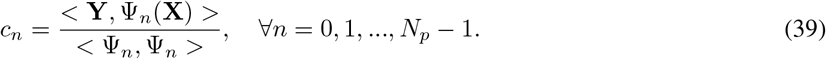

The equation above is obtained by taking the inner product of eq. (38) and Ψ_*n*_ and using the orthogonality of the basis. The coefficients are calculated with a pseudo-spectral projection method.

In particular, since we are using the uniform distribution *f*_**X**_ of the parameters, Ψ_*n*_ is based on Legendre polynomials, which are orthogonal with respect to the uniform distribution. These polynomials are obtained with a three-term recurrence formula,

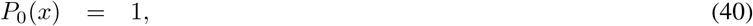

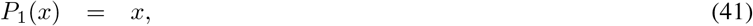

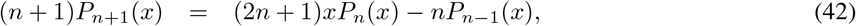

where *P*_*n*_(*x*), *n* ∈ ℕ, denotes the Legendre polynomials.

Once the simulations are complete, both an approximation of the model output and the calculated statistics of the QoIs based on the PCE are available.

Many statistical measures can then be obtained directly from the PCE representation, such as the mean and variance of the model response. The Sobol’ indices can also be computed based on the polynomial chaos decomposition of the model Sudret [2008] and are called PC-based Sobol’ indices.

## Results

### Simulating the healthy state

Parameters were tuned to simulate normal (healthy) conditions. The main characteristic is the existence of *γ* rhythm, as it is also observed in clinical studies Masimore et al. [2005], Kalenscher et al. [2010]. The current *I*_0_ (expressing dopamine functionality or cortical excitation) was set to 5 mA. Figure 4 depicts the overall neuronal activity of the striatum under these normal conditions.

**Fig 4:**
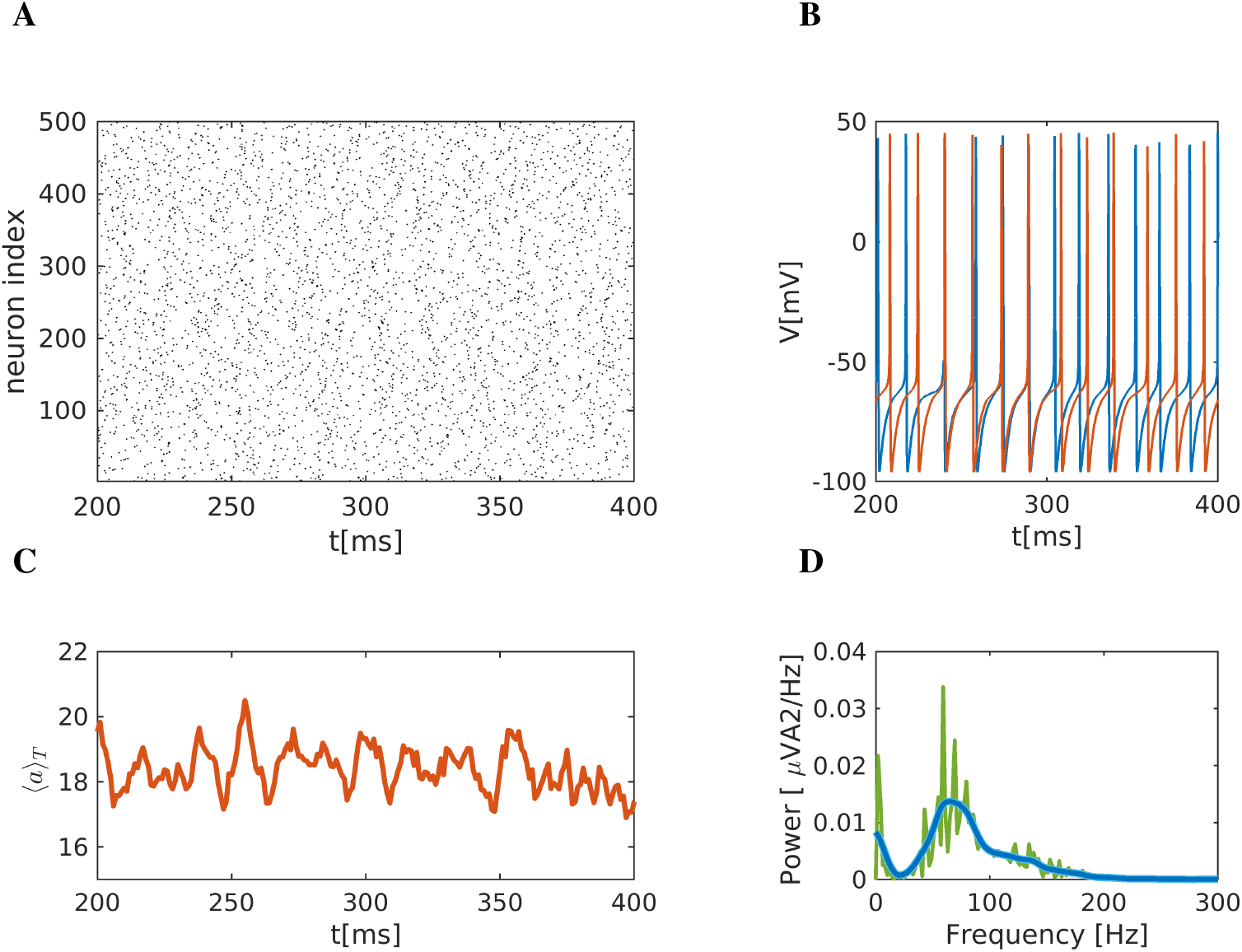
Representation of the striatal network dynamics under healthy conditions. **A** Raster plot representation. Black dots represent activated neurons (i.e. action potentials defined as transients passing *V* = − 15 mV to positive values) against time (in ms) and space (i.e. index of neuron of the nucleus). **B** Time series of two representative medium spiny neurons (MSN) of the striatum. **C** Mean activity of the striatal network. **D** Power spectrum of the mean membrane potential changes; see eq. (29), showing high activation in the *γ* band, i.e. at frequencies *>* 35 Hz. Green: High-resolution spectrum with a partitioning of 0.01 Hz. Blue: Smoothened curve using Gaussian function smoothing. In the high-resolution spectrum, the three main peaks are found at 59, 69 and 79 Hz, while in the smoothened one, the peak is at 65 Hz.

The raster plot (Fig. 4A) shows the activity of 500 randomly chosen neurons; the network apparently is not very much synchronised. Two representative MSN neurons are shown in Fig.4B. These neurons exhibit spiking activity with variable periods (i.e. non-constant period between two spikes), and some neurons appear to show brief intervals of synchronised activity, preceded and followed by non-synchronous firing. Such synchrony could either be due to transient common activation via network inputs (e.g. inhibition of fast-spiking neuron), or it could actually occur by chance with this tonic firing at a relatively high frequency. The mean network activity *r*_*t*_, i.e. eq. (28) as the macroscopic variable quantifying striatal activity, is depicted in Fig. 4C. It fluctuates around its mean value of 20 (action potentials/neuron within 10 ms). The Fourier analysis of the firing rate *r*_*T*_, in turn, is depicted in Fig. 4D. The main characteristic of the power spectrum is a broad interval of *γ* band activity (*f >* 35 Hz) with a main peak at ≈ 65 Hz and a secondary smaller peak at ≈ 3 − 5 Hz (blue curve in Fig. 4D).

### Simulating abnormal low cortico-striatal activity

Consistent with the hypothesis that both OCD and depression are associated with a reduction in dopaminergic (tegmental) and glutamatergic (cortical) activity Obeso and Lanciego [2011], Albin et al. [1995], Caravaggio et al. [2016], Stephens et al. [2005], we model a low excitation tone from the cortex to the striatum Stephens et al. [2005] which is referred in this paper as the “abnormal condition”. For this purpose, we reduce the excitation current from *I*_0_ = 5 mA to *I*_0_ = 1.5 mA, which significantly changes the behaviour of the system. Figure 5 shows the overall network dynamics under these abnormal conditions. The raster plot in Fig. 5A depicts the activity of 500 randomly chosen neurons. The main emergent characteristic is weak network activity, which is sparse and decreased compared to healthy activity. Two representative neurons are shown in Fig. 4B. The traces of membrane potential confirm the abnormally low activity, with long intervals of neuronal silence (no spiking activity). The mean network activity ⟨*a*⟩ _*T*_, i.e. eq. (28), is depicted in Fig. 4C. This mean network activity also reflects its reduced activity, which now fluctuates around action potentials 2 and 3 action potentials/neuron within 10 ms. Lastly, the Fourier analysis of the mean network activity ⟨*a*⟩ _*T*_ also captures this change; see Fig. 4D. The power spectrum is now shifted towards low frequencies with a main peak ≈ 4 Hz indicating a *θ* rhythm, a secondary peak at 22 Hz, i.e. *β* band, and a small peak at around 55 Hz.

**Fig 5:**
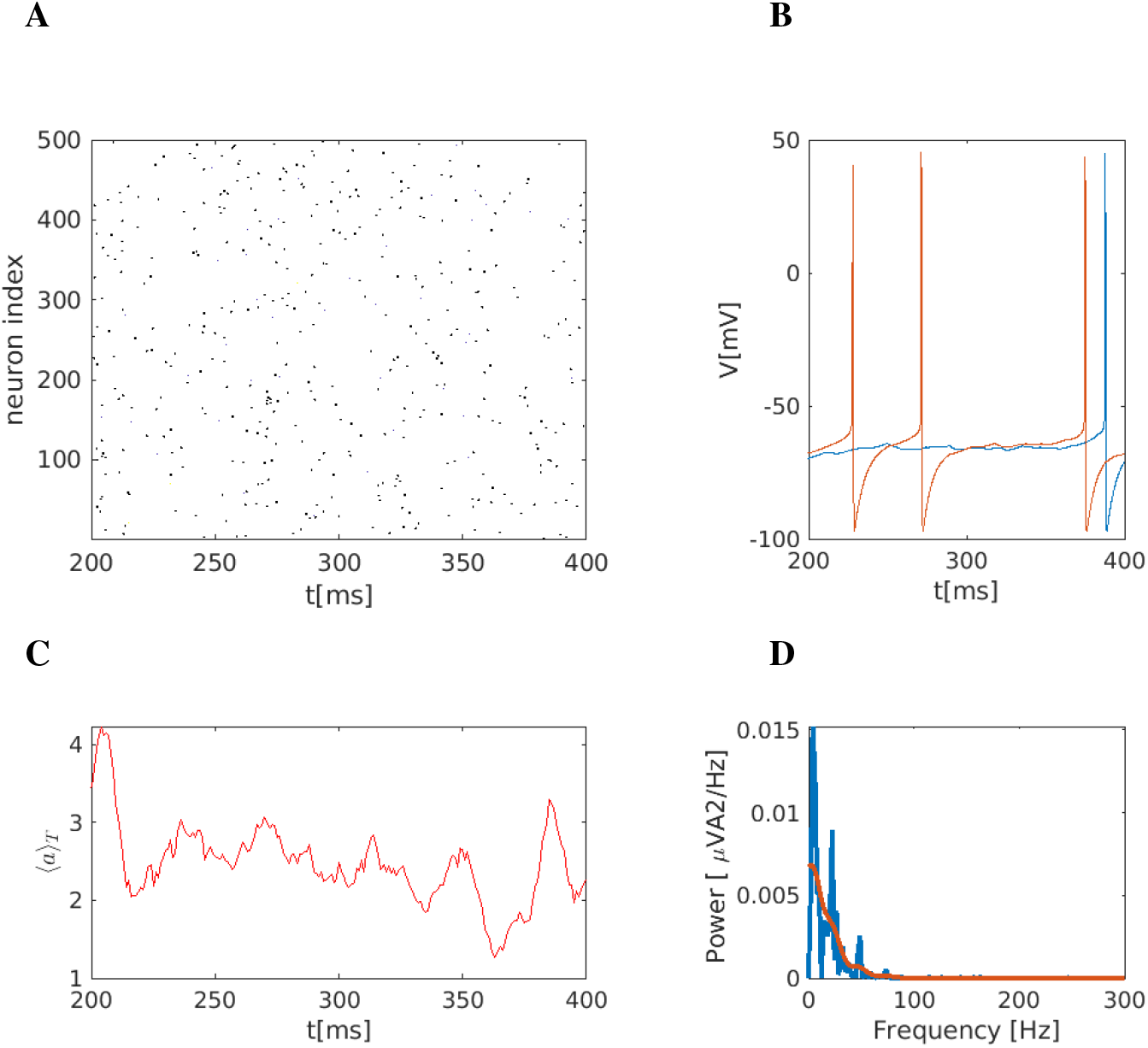
Representation of the striatal network dynamics under abnormal state at low external activation. **A** Raster plot representation. Black dots represent activated neurons (i.e. action potentials defined as transients passing *V* = − 15 mV to positive values) against time (in ms) and space (i.e. index of neuron of the nucleus). Compared to the healthy state, the raster plot shows very sparse activity. **B** Time series of two representative medium spiny neurons (MSN) of the striatum. **C** The mean activity rate of the striatal network shows abnormally low activity compared to a healthy state. **D** Power spectrum of the mean membrane potential changes; see eq. (29). Three dominant peaks emerge at ≈ 4, 22, and 55 Hz.

### Simulating abnormal state using optimal DBS parameters

In this section, we present the optimisation results for network dynamics based on the minimisation of (a) the differences of mean network activity under healthy and abnormal conditions, i.e. when minimizing the objective function eq. (30), (b) of network rhythmicity (as determined by the spectrum of the mean membrane activity on the network which is correlated to local field potentials (LFP)) and mean network activity and of (c) mean network activity of the dorsal striatum only (e.g. considering a specific targeted area which is known to be involved in neurpsychiatric disorders). To model the influence of DBS on a network in an abnormal state, the network structure and the model parameters were kept at the abnormal striatum state. Based on the deviation from the healthy state, we propose a DBS protocol that provides the therapeutic pattern for abnormal striatal network activity.

### Optimised DBS parameters with respect to the mean network activity

The optimal values for the position, frequency and amplitude of DBS were found to be *r* = (*x*_0_, *y*_0_, *z*_0_, *A*_DBS_, *f*) = (− 11.39, 4.21, 15.11, 195.02, 99.73). The optimal position for DBS, together with a network snapshot, is depicted in Fig. 6A. The raster plot (Fig. 6B) shows the strongly synchronised activity of the network due to DBS. The mean network activity is depicted in Fig. 6C, jointly with healthy, pathological, and DBS at the initial position, for comparison reasons. Clearly, the mean network activity resulting from optimised (blue line) DBS is in very good agreement with the mean network activity found in the healthy state (red one). The spectrum (Fourier analysis) of the mean activity 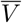 is shown in Fig.4D. The power spectrum shows high peaks at ≈ 100 Hz and ≈ 200 Hz as a result of high-frequency DBS. Finally, firing patterns of two representative neurons are depicted in Fig.4E. The neurons show activity restored close to healthy conditions.

**Fig 6:**
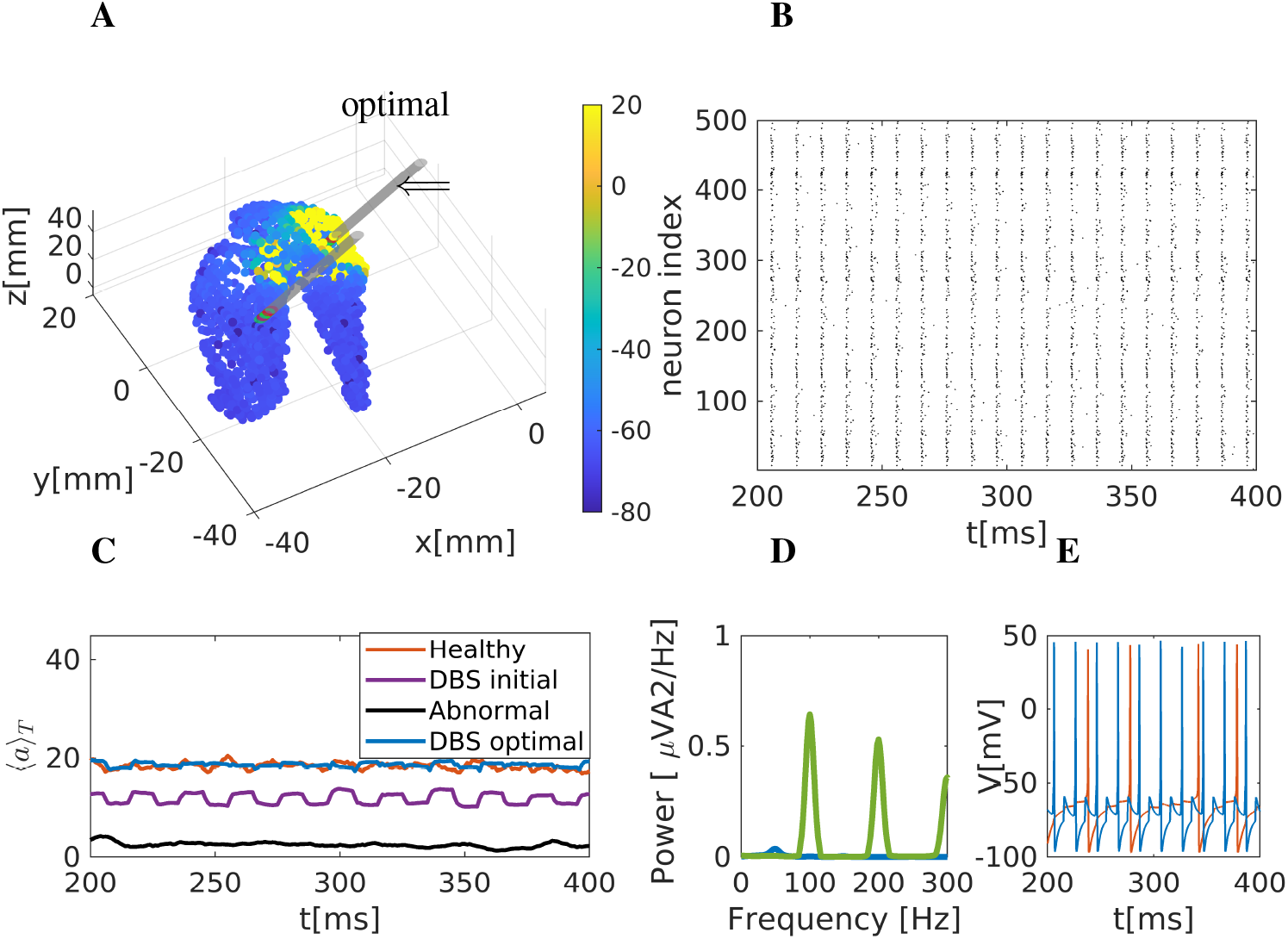
Optimised DBS activity on striatal network. **A** Snapshot of striatal activity during DBS. The colour code depicts the predicted mean membrane potential of neurons affected by the stimulation (in mV). Two electrodes are included: one corresponds to the optimal position, marked with an arrow, while the second is the initial position at the beginning of the optimisation process. **B** Raster plot representation. Black dots represent activated neurons. Due to the stimulation, the activity is highly synchronised. **C** Mean network activation of the striatal network. The thick blue line (mean network activity) results from stimulating at optimal DBS position. For comparison, we also show mean activities under healthy, pathological, and DBS conditions with stimulation in the initial, non-optimised position. **D** Power spectrum of the firing rate. **E** Two representative neurons were simulated using a stimulation set at optimal DBS conditions.

### Optimised DBS parameters with respect to the network rhythmicity and mean network activity

We next used a third objective function, *obj*_3_, which optimises stimulation with respect to the network phase and the mean membrane activity. The optimal values for the position, frequency and amplitude were found to be *r* = (*x*_0_, *y*_0_, *z*_0_, *A*_DBS_, *f*) = (− 6.46, 4.87, − 0.75, 367.89, 109.19). The optimal stimulation position together with a network snapshot is depicted in Fig. 7A, The raster plot (Fig. 7B) shows a strong periodic activity at 110 Hz due to the high-amplitude, high-frequency DBS, similar to the previous optimisation case. The mean network activity is depicted in Fig. 7C, jointly with mean network activities under healthy and pathological states, as well as DBS conditions with DBS at the initial position. The resulting firing rate (using *obj*_3_) shows a spike-like activity mainly related to the high-amplitude DBS stimulation, i.e. *A*_DBS_ = 367.89. The spectrum (Fourier analysis) of the mean membrane potential 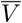 is shown in Fig.7D. The power spectrum now shows the highest peak at ≈ 110 Hz and a secondary peak at 220 Hz, i.e. as expected in the range of the stimulation frequency and its harmonics. Finally, the firing activity of two representative neurons is depicted in Fig.7E. Interestingly, one neuron follows the DBS high frequency firing at 110 Hz, while the other is actually unaffected and firing at low frequency, showing that not all neurons are recruited into the stimulation.

**Fig 7:**
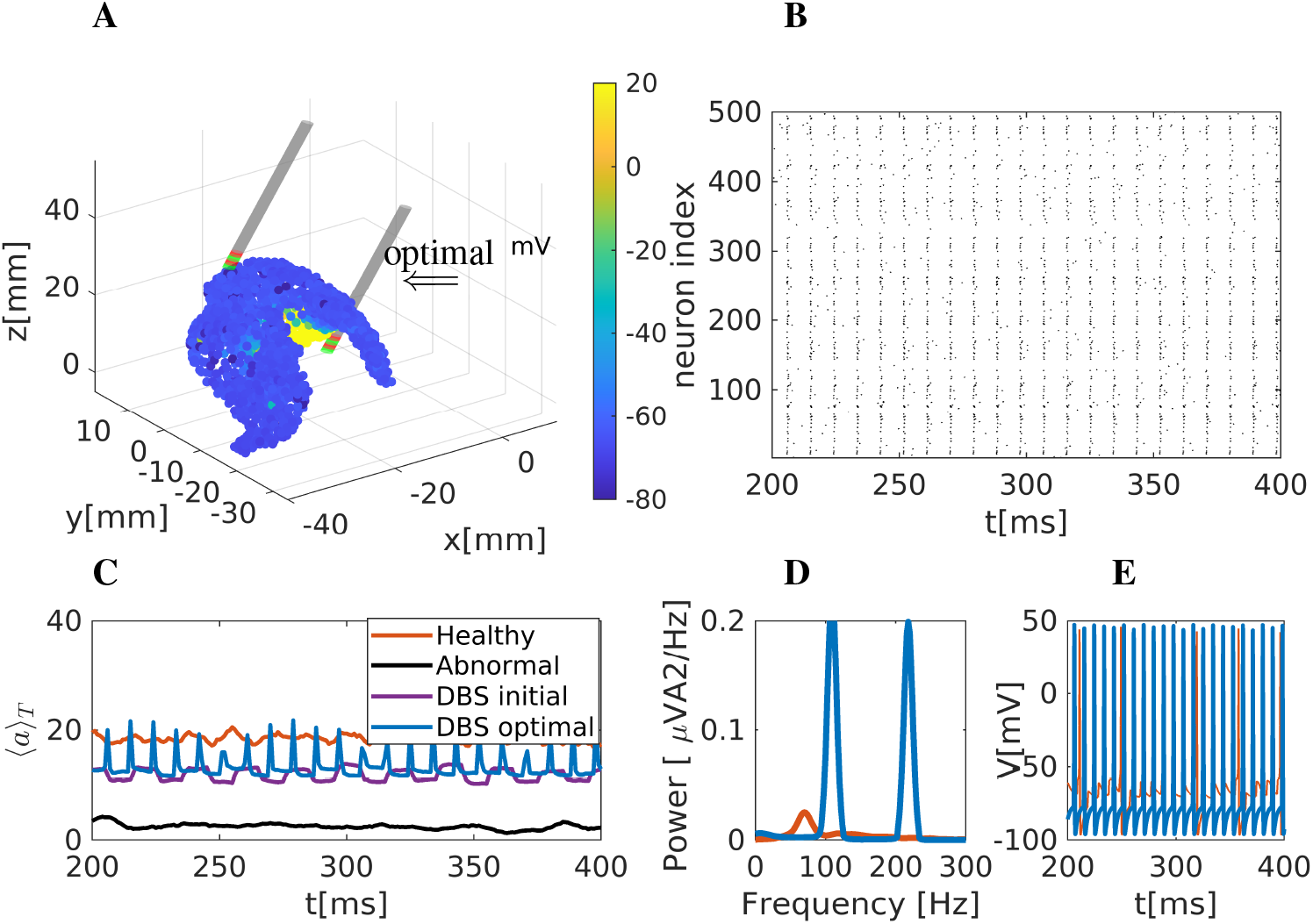
Optimised DBS with respect to the phase and the firing rate of the striatum network. **A** Snapshot of striatal activity during DBS. The colour code depicts the predicted mean membrane potential of neurons affected by the stimulation (in mV). Two electrodes are included: one corresponds to the optimal position, marked with an arrow, while the second is the initial position at the beginning of the optimisation process. **B** Raster plot representation. Black dots represent activated neurons. Due to the stimulation, the activity is highly synchronised. **C** Mean network activation of the striatal network. Blue thick line (mean network activity) resulting from stimulating at optimal DBS position. For comparison reasons, we also show mean activities under healthy, pathological, and DBS conditions with stimulation in the initial, non-optimised position. **D** Power spectrum of the firing rate. **E** Two representative neurons were simulated using a stimulation set at optimal DBS conditions.

### Optimised DBS with respect to the activity of a specific targeted area (dorsal striatum)

In our network modularity analysis, a partitioning of the striatum into six major areas emerges (see Fig.3), of which the dorsal striatum is the most strongly affected by stimulation. For this reason, in the next step, we again optimised with respect to the firing rate, i.e. using *obj*_1_. This time, however, we only restricted the analysis to area 2 in Fig.3. Under these conditions, the optimal values for the position, frequency and amplitude were found to be *r* = (*x*_0_, *y*_0_, *z*_0_, *A*_DBS_, *f*) = (− 11.43, 4.26, 20.81, 200.16, 98.94). The optimal position together with a network snapshot is depicted in Fig. 8A, The raster plot (Fig. 8B) shows activity around 110 Hz due to the DBS effect. The mean network activity is depicted in Fig. 8C, again with the results from healthy and pathological (abnormal) conditions. The optimised condition (with respect to the phase, blue thick line) approximates the healthy state (red line). The spectrum (Fourier analysis) of the neuronal firing rates *r*_*T*_ is shown in Fig.8D, again showing 110 Hz and harmonics as peaks, as expected. Finally, two representative neurons are depicted in Fig.8E. The neurons show restored activity resembling the healthy state.

**Fig 8:**
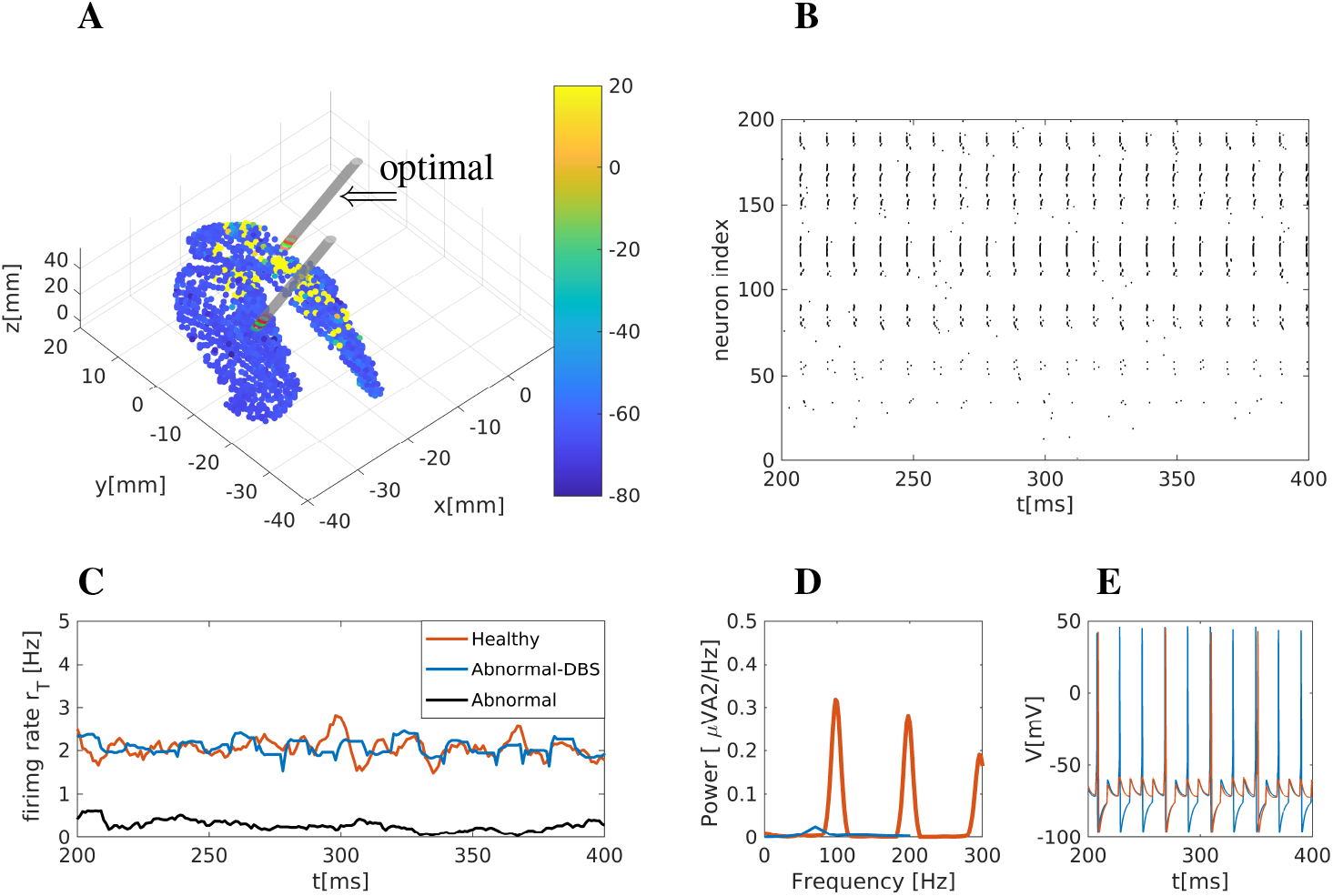
Optimised DBS activity for a specific targeted striatum area. **A** Snapshot of the striatum activity during DBS. Colour coding is according to the membrane potential (in mV). Two electrodes are included. One corresponds to the optimal position, marked with an arrow, while the second displays the initial position. **B** Raster plot representation. Black dots represent activated neurons, and the activity is synchronised. **C** Mean activity of the striatal network. Blue thick line (firing rate) resulting under optimal DBS positioning conditions. We also compare mean activities under healthy, pathological, and DBS conditions with stimulation in the initial, non-optimised position. **D** Power spectrum of the firing rate. **E** Two representative neurons were simulated using a stimulation set at optimal DBS conditions.

### Sensitivity analysis using Polynomial Chaos Expansion

We have performed a sensitivity analysis using polynomial chaos expansion (PCE) to study the sensitivity of the parameters involved in our model. For this analysis, we consider the parameters *I*_0_, cortical excitation current and *g*_*MM*_, i.e. conductance between MSN neurons, defined in the intervals [1.5, 5.0] and [0.0, 0.1], respectively. The parameter values are sampled following a uniform distribution. *n*_*s*_ = (*p* + 1)^*d*^ gives the number of samples, with *p* = 3 the order of the polynomial chaos expansion and *d* = 2 the number of parameters. We have 16 samples to run the model, each running in approximately 35 minutes. The macroscopic quantity of interest (QoI) is the firing rate of neurons. Implementing the uncertainty quantification routine is based on the EasyVVUQ library Richardson et al. [2020] in Python. The analysis approach is non-intrusive, i.e., the model is considered a black box. Figure 9 presents the first-order and the higher-order Sobol’ indices obtained for the described configuration. As pointed out earlier in the methods section, Sobol’ indices correlate with the relative impact of any given parameter on the uncertainty in the output parameters. The simulation was performed for a time window from 200 ms to 400 ms. First, these indices were calculated, letting both parameters interfere with each other, and in a second step, the simulation was run for each parameter (*I*_0_ and *g*_*MM*_) independently. When both parameters can interfere with each other, it is visible that *g*_*MM*_ has more impact than *I*_0_ (Fig. 9A). When analysing the impact independently, *I*_0_ remains below values of 0.4 (Fig. 9B), while *g*_*MM*_ has a higher impact with an average value of 0.6 (Fig. 9C). Thus, although *I*_0_ influences the firing rate, it is less significant than the variation of *g*_*MM*_. While fluctuating over time, the first-order index *g*_*MM*_ has an average value of 0.6, showing considerable importance in the model response (Fig. 9). One interpretation is that while cortical input, defined by *I*_0_, generates the initial drive, *g*_*MM*_ considerably boosts these cortical inputs. In other words, the internal conductance of MSN neurons plays a significant role in the network dynamics. It is conceivable that any pathological reduction of MSN activity or even dendritic remodelling in these cells will reduce the effect of glutamatergic and dopaminergic inputs onto the striatum and lead to abnormal striatal activity. Our results on sensitivity analysis also support this hypothesis.

**Fig 9:**
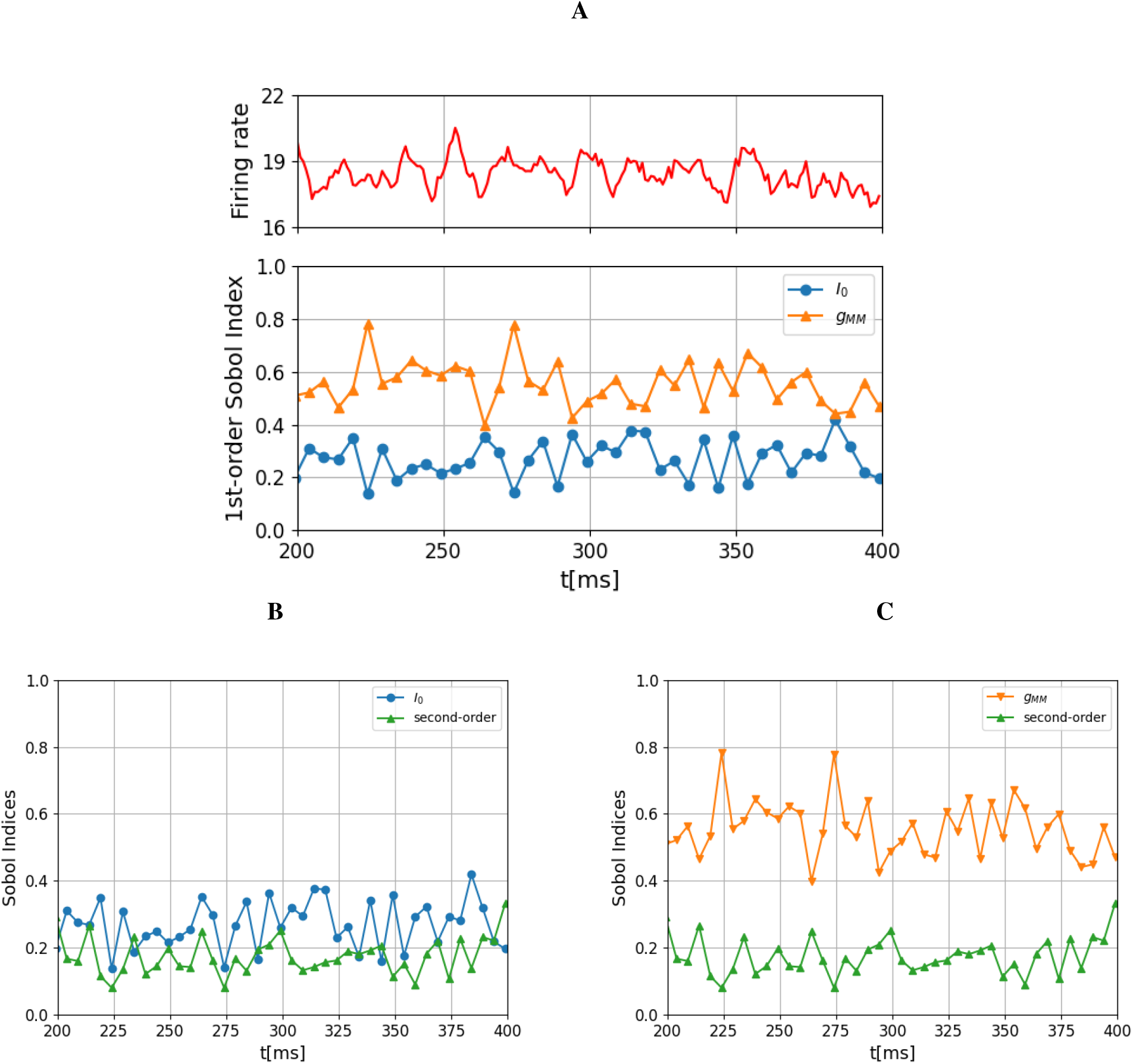
First- and higher-order Sobol indices over time in the interval [200 ms, 400 ms]. **A** On top, the firing rate is shown. Below, For the first order Sobol’ indices of the two parameters, cortical excitation current (*I*_0_) and conductance between MSN neurons (*g*_*MM*_), are displayed. The conductance between MSN neurons (*g*_*MM*_) plays a dominant role in the firing rate.**B** Individual contribution of *I*_0_ and the second order Sobol index. The latter indicates the joint contribution of both *I*_0_ and *g*_*MM*_ to the output variance due to interactions between them. **C** This figure shows the individual contribution of *g*_*MM*_ and the second order index. The range of the second-order Sobol index suggests that the interactions between these parameters contribute to the output variance, but not as much as the individual contributions of *I*_0_ and *g*_*MM*_ separately. Therefore, the conductance of MSN neurons influences the firing rate.

## Discussion

We developed a biophysical network model for the striatum to explore the relationship between anatomical structure and neural activity, allowing us to calculate optimal DBS parameters based on spatio-temporal patterns produced by the model. The network construction was based on (a) an FDA-approved state-of-the-art human atlas Iacono et al. [2015] (extracting coordinates for the striatal neurons), (b) on modified Hodgkin-Huxley equations for medium spiny neurons (MSN) and fast-spiking neurons (FSN) Chartove et al. [2020], Hodgkin and Huxley [1952], and (c) on complex network structures for neuronal connectivity Netoff et al. [2004], Berman et al. [2016], She et al. [2016], Bassett and Bullmore [2006, 2017], Fang et al. [2017], de Santos-Sierra et al. [2014].

Depending on the model parameters, the network produced three spatiotemporal patterns, i.e. healthy, abnormal/pathological (presumably mirroring the situation in psychiatric disorders such as OCD and depression; Caravaggio et al. [2016]), and DBS conditions. Simulating healthy conditions, the neuronal model produces macroscopic network activity with two main spectrum components, one peak on the *θ* rhythm (around 5 Hz) and the main peak at *γ* frequency band (60 Hz, see Fig. 4). This *γ* band activity is also observed in animals (rats) clinical studies of striatum Masimore et al. [2005], Kalenscher et al. [2010] during the movement initiation or for motivated behaviour and reward processing Masimore et al. [2005], Kalenscher et al. [2010]

The model has shown reduced striatal activity by changing the conditions, specifically by reducing the cortico-striatal excitatory tone. This is also depicted in the spectrum diagram (see, Fig. 5), where, in contrast to healthy conditions, the highest peak appears at 4 Hz (*θ* band), while a secondary peak exists at 22 Hz (*β* band) and a third one around 48 Hz. The reduced excitatory tone is explainable by a possible reduction of dopaminergic and glutamatergic inputs presumably occurring in neuropsychiatric conditions: Caravaggio et al. Caravaggio et al. [2016] showed that chronic dopamine depletion (*>* 4 months) produces decreases in striatal glutamate (consistent with the classical model of the basal ganglia). Dopamine reduction, in turn, has been observed in Major Depressive Disorder Pizzagalli et al. [2019]. Furthermore, decreased functional connectivity (or decreased excitatory glutamatergic tone) between the sensorimotor cortex and dorsolateral striatum, and between dorsomedial striatum and dorsolateral prefrontal cortex has also been observed in PD patients Mi et al. [2021], Rao et al. [2010]. Patients with obsessive-compulsive disorder, in turn, when carrying out a cognitive task, showed decreased responsiveness in the right medial and lateral orbitofrontal cortices, as well as in the right caudate nucleus (meaning to say, the cranial part of the striatum) when compared to controls Remijnse et al. [2006].

The proposed optimisation process was based on (a) the definition of objective functions (and thus possible biomarkers) that measure (or characterise) the network activity patterns and (b) the deviation of these biomarkers from the healthy state (the optimisation aiming to minimise this deviation). Our method estimates the optimal DBS parameters (including the following parameters: position, amplitude and frequency of the electrical signal) by a repetitive process, eventually aiming to restore or at least approximate healthy neuronal activity of the striatal network.

The first optimisation protocol relied on the striatal network’s mean activity (⟨*a*⟩). Following this optimisation procedure, the parameter ⟨*a*⟩ approximated neural activity parameters in the healthy state to a large degree. The optimal DBS parameters were found to be *r* = (*x*_0_, *y*_0_, *z*_0_, *A*_DBS_, *f*) = (−9.62, 2.13, 11.22, 251.85, 99.54). The second optimisation approach was based on the combination of the network rhythmicity and mean network activity. The optimal parameter in that case was found to be *r* = (*x*_0_, *y*_0_, *z*_0_, *A*_DBS_, *f*) = (− 6.46, 4.87, − 0.75, 367.89, 109.19). Interestingly, in primates, high (at 200 Hz, rather than low at 20 Hz) frequency stimulation appears to improve cognitive function even under healthy conditions when delivered to the striatum Williams and Eskandar [2006]. The third optimisation approach was based on the structural connectivity of the network. Specifically, we utilised modularity analysis and the resulting partitioning of the striatum (see Fig.3). We guided our stimulation to the dorsal striatum. We restricted the analysis to area 2 in Fig.3. A final optimisation approach (depicted in the supplementary) was also implemented, based only on Fourier analysis of neuronal action potentials, shown in Fig. 10. where 64 Hz DBS restored mean firing frequencies from *θ* in the abnormal case, to *γ* close to the healthy state.

**Fig 10:**
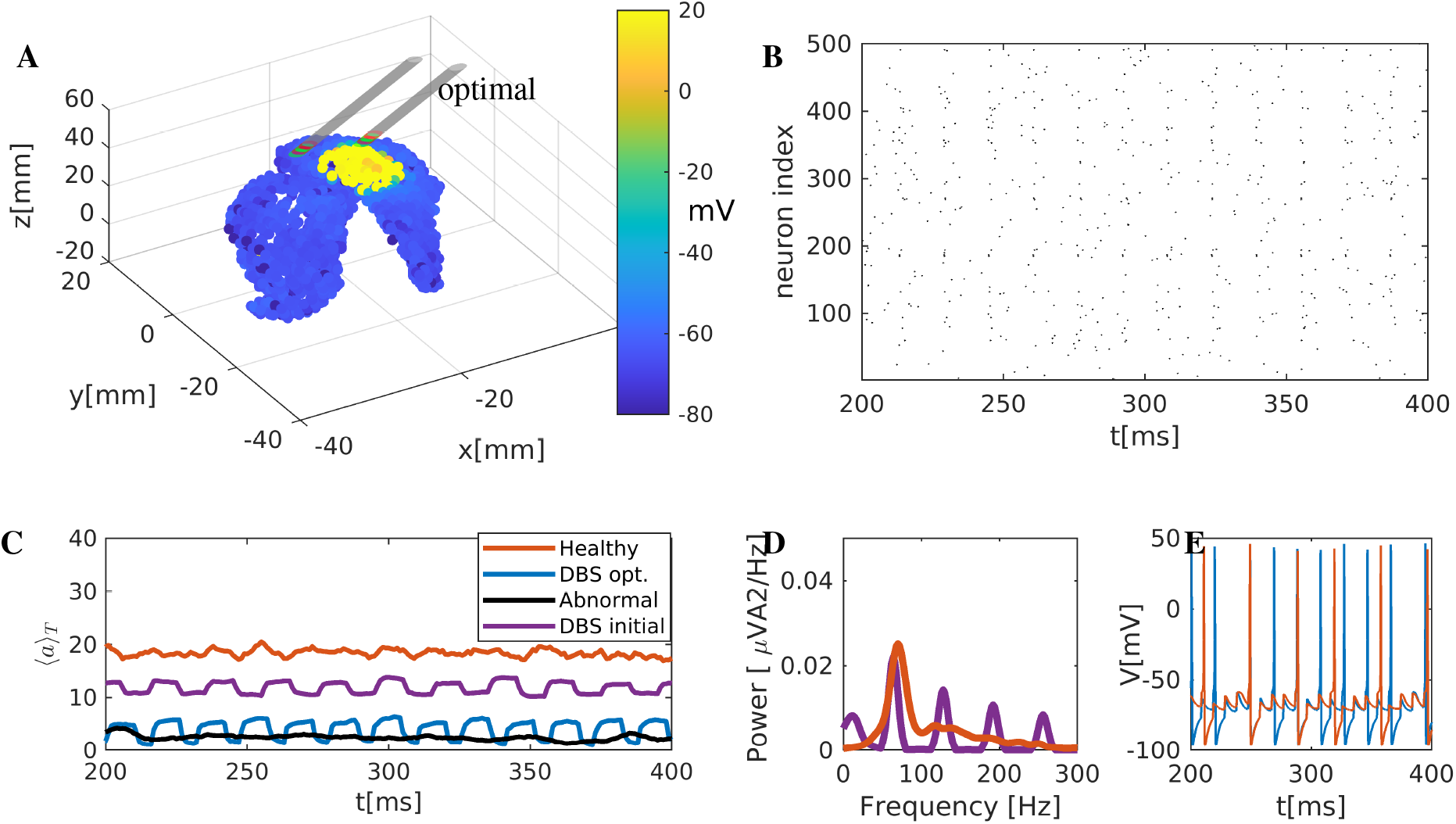
Optimised DBS with respect to the network rhythmicity. **A** Snapshot of the striatum activity during DBS. Colour coding is according to the membrane potential (in mV). Two electrodes are included; one corresponds to the optimal position, marked with an arrow, while the second marks the initial position. **B** Raster plot representation. Black dots represent activated neurons, and the activity is synchronised. **C** Firing rates of the striatum network. Blue thick line (firing rate) resulting from the optimal DBS position. For comparison reasons, we include the healthy, pathological and DBS on a refereed position. **D** Power spectrum of the firing rate. **E** Two representative neurons were simulated using optimal DBS conditions.

Different methods for finding optimal targeting positions for striatal DBS have been reported in the literature Sheth et al. [2022], Makris et al. [2016], Li et al. [2020], van Westen et al. [2021]. A clinical protocol is described in van Westen et al. [2021], where critical parameters, e.g. the amount of current that is applied, the number of electrical pulses per second (frequency), the duration of these pulses (pulse width), and the amplitude of these pulses (similar to our parameters estimation), are tuned gradually depending on the positive performance of the patient. Other methods use hybrid approaches, combining clinical and computational methods, usually by correlating the activation of fibre bundles (calculating the volume of tissue activated) with patients’ optimal clinical response Sheth et al. [2022], Makris et al. [2016], Li et al. [2020]. As proposed in Li et al. [2020], axons (fibre) activation modulates neuronal network activity responsible for clinical improvement. However, fibre tracts and the volume of tissue activated do not provide any information on the reaction of the neuronal network (i.e. how the tissue activation or bundle activation modulates the neural activity). Only a recent publicationSheth et al. [2022] studied the response of specific brain networks using an indirect biomarker, the intracranial electroencephalogram (EEG). In this paper, the authors identified hubs of critical white matter pathways (using tractography) connecting cortical and subcortical network regions relevant to the expression of depressive symptoms Sheth et al. [2022]. Our approximation constitutes a different, supplementary approach to the aforementioned studies based on cortical-striatal network activation. These two network parts are intricately entangledPeters et al. [2021]; according to these authors, the relationship between neural activity in the cortex and striatum is “spatiotemporally precise, topographic, causal and invariant to behaviour, supporting, thus, a causal role of cortical inputs in driving the striatum” Peters et al. [2021]. Finally, for completeness, we refer to multiple surgical targets for treating obsessive-compulsive disorder with deep brain stimulation (DBS) Moser et al. [2011]: ant. cingulate cortex (−7.9,27.2,-7) and (−6.5,1.6,-4). Targets in Nucleus Accuben (−7.5, 10.8, -5) and in ventral striatum / ventral capsule (−8.4, 3.5, -1), (−7.5, 15.3, -5).

Our study constitutes a computational approximation of the complex striatal network with assumptions and limitations. Regarding the assumptions, we used two types of neurons with simplified equations (see Chartove et al. [2020]). Furthermore, internal connectivity in the nuclei was assumed to take the form of small-world complex structures. This novel approach in basal ganglia modelling has a reasonable justification in previous publications, both modelling and experimental Netoff et al. [2004], Berman et al. [2016], She et al. [2016], Bassett and Bullmore [2006, 2017], Fang et al. [2017], de Santos-Sierra et al. [2014]. As a limitation of the model, the exact structure of the connectivity on this microscopic level is unknown. Hence, how this can be modelled in the future remains to be clarified.

From a future modelling perspective, one important step forward will be the integration-connection of the striatal model with cortical areas, as well as with the other basal ganglia and thalamic nuclei. The new augmented model should contain synaptic plasticity effects (potentiation, depression), both known to be present at corticostriatal synapses, which strongly depend on the activation of dopamine receptors Calabresi et al. [2007]. An integrative model will give new insight into possible mechanisms of DBS.

## Acknowledgments

The structural MRI data were provided [single subject data] by the Human Connectome Project, WU-Minn Consortium (Principal Investigators: David Van Essen and Kamil Ugurbil; 1U54MH091657) funded by the 16 NIH Institutes and Centers that support the NIH Blueprint for Neuroscience Research; and by the McDonnell Center for Systems Neuroscience at Washington University.

## Funding

This work was funded by the Deutsche Forschungsgemeinschaft (DFG, German Research Foundation) – SFB 1270/2 - 299150580 - Collaborative Research Centre ELAINE.

## Author contributions statement

KS, RK, JS and RA contributed to the conceptualisation, methodology, model analysis and investigations. RA, JPP, and AKFG were involved in model simulations and investigations. RA, RK, JS, and UvR provided supervision throughout the project. KS, RA, RK, JS, SA, and UvR contributed to reviewing and editing the manuscript. All authors contributed to the article, writing of the original draft, and approved the submitted version.

## supporting material

### S1 File. optimisation with respect to the rhythmicity (Fourier spectrum) of the network

We use the second objective function based on the power spectrum of the mean membrane activity. The optimal DBS parameters will produce patterns according to the similarity of the DBS network rhythm compared to the healthy one. The optimal values for the position, frequency and amplitude were determined as *r* = (*x*_0_, *y*_0_, *z*_0_, *A*_DBS_, *f*) = (− 15.23, 3.37, 34.30, 253.74, 64.02). The optimal position, together with a network snapshot, is depicted in Fig. 10A. The raster plot (Fig. 10B) shows sparse activity with periodicity around 60Hz due to the DBS effect. The mean network activity is depicted in Fig. 10C, jointly with healthy, pathological DBS in a position close to the refereed for OCD for comparison reasons. The optimised (with respect to the phase, blue thick line) approximates the healthy state (black one). The spectrum (Fourier analysis) of the firing rate *r*_*T*_ is shown in Fig.4D. The power spectrum now shows the highest peaks at ≈ 55, 20 Hz and secondary peaks at 100 and 200 Hz. Finally, two representative neurons are depicted in Fig.4E. The neurons restore activity close to the healthy one.

## Notes

### Competing Interest Statement

The authors have declared no competing interest.

